# Comprehensive mapping of the Interaction of levodopa and iron metabolism in Parkinson’s disease

**DOI:** 10.1101/2024.09.13.612928

**Authors:** Jian Wang, Srinivasan Ekambaram, Xuemei Huang, Richard B. Mailman, Elizabeth A. Proctor, Nikolay V. Dokholyan

**Affiliations:** Department of Pharmacology, Penn State College of Medicine, Hershey, PA 17033, USA; Department of Neurology, PO 800394, UVA Health System, Charlottesville, VA 22908; Department of Neurology, Penn State Milton S. Hershey Medical Center; Department of Neurosurgery, Penn State Milton S. Hershey Medical Center and College of Medicine, Hershey, PA 17033, USA; Department of Biochemistry & Molecular Biology, Penn State College of Medicine, Hershey, PA 17033, USA; Department of Chemistry, Pennsylvania State University, University Park, PA 16802, USA; Department of Biomedical Engineering, Pennsylvania State University, University Park, PA 16802, USA; Department of Engineering Science & Mechanics, Pennsylvania State University, University Park, PA 16802, USA

**Author notes:** These authors contributed equally to this work (Jian Wang, Srinivasan Ekambaram).

## Abstract

Levodopa remains the primary treatment for Parkinson’s disease (PD), yet its long-term use has been associated with iron accumulation in the brain, a phenomenon linked to neurodegeneration. We utilize deep machine learning to determine plausible molecular mechanisms that may underlie the effects of levodopa on iron metabolism. Using the DRIFT platform, we performed a proteome-wide target identification of levodopa and uncovered significant interactions potentially involved in cellular iron transport. Pathway analysis revealed that levodopa may influence critical iron-related pathways, including the response of EIF2AK1 to heme deficiency, heme signaling, and ABC-family protein-mediated transport. These findings suggest that levodopa may contribute to iron dysregulation in PD by interacting with iron transporters and modulating iron-related pathways. Because levodopa is used at relatively high doses in PD, our findings provide new insight into secondary effects unrelated to being a precursor of dopamine. This highlights the need for careful consideration of its effects on iron metabolism as a consequence of use in the long-term management of PD. Further experimental validation is required to confirm these interactions, and also to explore potential strategies to mitigate iron-related side effects while preserving therapeutic efficacy.

## Introduction

Levodopa has been the gold standard in the treatment of Parkinson’s disease (PD) for more than a half-century^1^. It overcomes the effects of dopamine neuron degeneration in the substantia nigra by increasing dopamine production, thus alleviating the motor symptoms of PD^2^ and significantly improving quality of life of patients. While there is still no approved drug that can substitute for levodopa, there has been significant controversy about potential undesired actions. One of the mechanisms for these undesired effects was the reported accumulation of iron in the brain of PD patients. It was proposed that the high concentrations of Fe2+^3^ and the presence of levodopa and its downstream products could generate reactive oxygen species (ROS)^4^. These ROS then would cause neurodegeneration^5^.

This levodopa-iron toxicity hypothesis, coupled with levodopa-related motor fluctuation and dyskinesia, led to a change in the PD standard-of-care for early disease (i.e., delayed start of levodopa; see reviews in ^6–9^). “Dopamine agonists” (D2-selective drugs) were used for early therapy to avoid the “toxic” effects of levodopa until patients could no longer tolerate inadequate control of symptoms. It was recognized, however, that a controlled clinical trial was needed for evidence, and this led to the landmark **ELLDOPA** study (**E**arlier versus **L**ater **L**evo**DOPA** Therapy in PD)^10^. Contrary to the levodopa-toxicity hypothesis, ELLDOPA found that levodopa had clinically beneficial effects on patients rather than the predicted negative effects^9^. Another major study, **LEAP** (***L****evodopa in* ***Ea****rly PD)*^11,12^ also found no evidence for the predicted toxicity of levodopa. These findings returned the standard of care to early use of levodopa.

There remained, however, a conundrum. *In vivo* radio imaging measuring the density of striatal dopamine transporters (DAT) has been developed as a biomarker of the integrity of dopaminergic nerve terminals^13^. This became an important dependent variable in the ELLDOPA study. While neither ELLDOPA nor LEAP found functional damage in the levodopa-treated arm, there was a significantly lower DAT density in this group. Patient well-being was, however, more important than a biomarker, so this observation remained stagnant.

Studies related to the accumulation of iron continued, aided by the ability to use susceptibility MRI to estimate iron accurately *in vivo*. The original postmortem findings of increased iron were widely confirmed using MRI and often were associated with the state of disease assessed by side effects or time since diagnosis^14–22^. Our interest in this question became acute when we reported that while iron accumulation occurred as PD disease progressed, at the time of initial diagnosis (i.e., before initiation of levodopa therapy) the iron concentration was slightly lower than in age-matched controls^23^. Thus, there was no accumulation of iron despite the fact that in these newly diagnosed patients, it is clearly established that half or more of dopamine neurons and terminals have already disappeared. This caused us to speculate that both sides of the original controversy were right: Restoration of function with levodopa is essential for brain health (explaining the ELLDOPA and LEAP) clinical findings, but that levodopa may also be having concomitant toxicity that increases the rate of disease progression. Even though levodopa is essential for treatment, determining if it also causes toxicity is essential since current and developing technology may be able to decrease or avoid this problem while still giving patients optimal therapy.

The exact mechanisms by which levodopa might influence and/or interact with iron metabolism remain unclear. Traditional research methods have provided limited insights into this complex interaction, highlighting the need for novel approaches that can unravel the molecular underpinnings of levodopa’s effects on iron homeostasis. In this context, computational tools offer a powerful means of exploring drug-target interactions on a proteome-wide scale, enabling the identification of potential molecular targets and pathways that might be influenced by levodopa.

To this end, we have employed DRIFT^24^, an advanced deep learning-driven platform designed for rapid and comprehensive mapping of chemical compounds to their corresponding protein targets. The ability of DRIFT to predict small molecule-protein interactions across the entire proteome provides a unique opportunity to identify previously unrecognized targets of levodopa, particularly those involved in iron metabolism. Additionally, we performed pathway analyses to find pathways related to iron regulation that might be impacted by levodopa.

Our findings reveal a complex network of interactions between levodopa and various solute carrier transporters, particularly within the solute carrier organic anion transporter family (SLCO) and solute carrier family 22 (organic cation transporters, OCTs). These transporters play critical roles in the transport of iron and other molecules across cellular membranes, suggesting that levodopa may influence iron homeostasis through its interaction with these proteins. Furthermore, we identified key pathways involved in iron metabolism that may be modulated by levodopa, providing new insights into the potential mechanisms underlying its effects on iron levels in the brain.

## Results

### Similar Compounds of Levodopa Identified by DRIFT

We used DRIFT to first identify compounds similar to levodopa (Figure 1). The identified compounds share significant structural similarities with levodopa, primarily in their core phenolic ring and side chain configurations (Figure 2A). Levodopa itself contains a catechol group (a phenyl ring with two *o*-hydroxyl groups) attached to an alkyl side chain. This catechol moiety is a common denominator in engaging similar biological interactions. Melevodopa and droxidopa both retain the catechol group, though they feature different substituents on their amino acid chains, with melevodopa incorporating additional hydroxyl groups and an ester linkage, and droxidopa possessing an additional hydroxyl group on the side chain, contributing to variations in solubility and reactivity. Racemetirosine, metirosine, and tyrosine closely resemble levodopa with a similar side chain, maintaining the basic amino acid backbone but differing in the exact positioning and stereochemistry of the amino and carboxyl groups. Etilevodopa and methyldopa also retain this structural framework but introduce ester groups, enhancing lipophilicity and potentially altering metabolic stability. Carbidopa, while more structurally distinct, still preserves the catechol group, but introduces a hydrazine moiety that is key to its inhibition of aromatic amino acid decarboxylase, thus decreasing the peripheral conversion of levodopa to dopamine.

**Figure 1.**
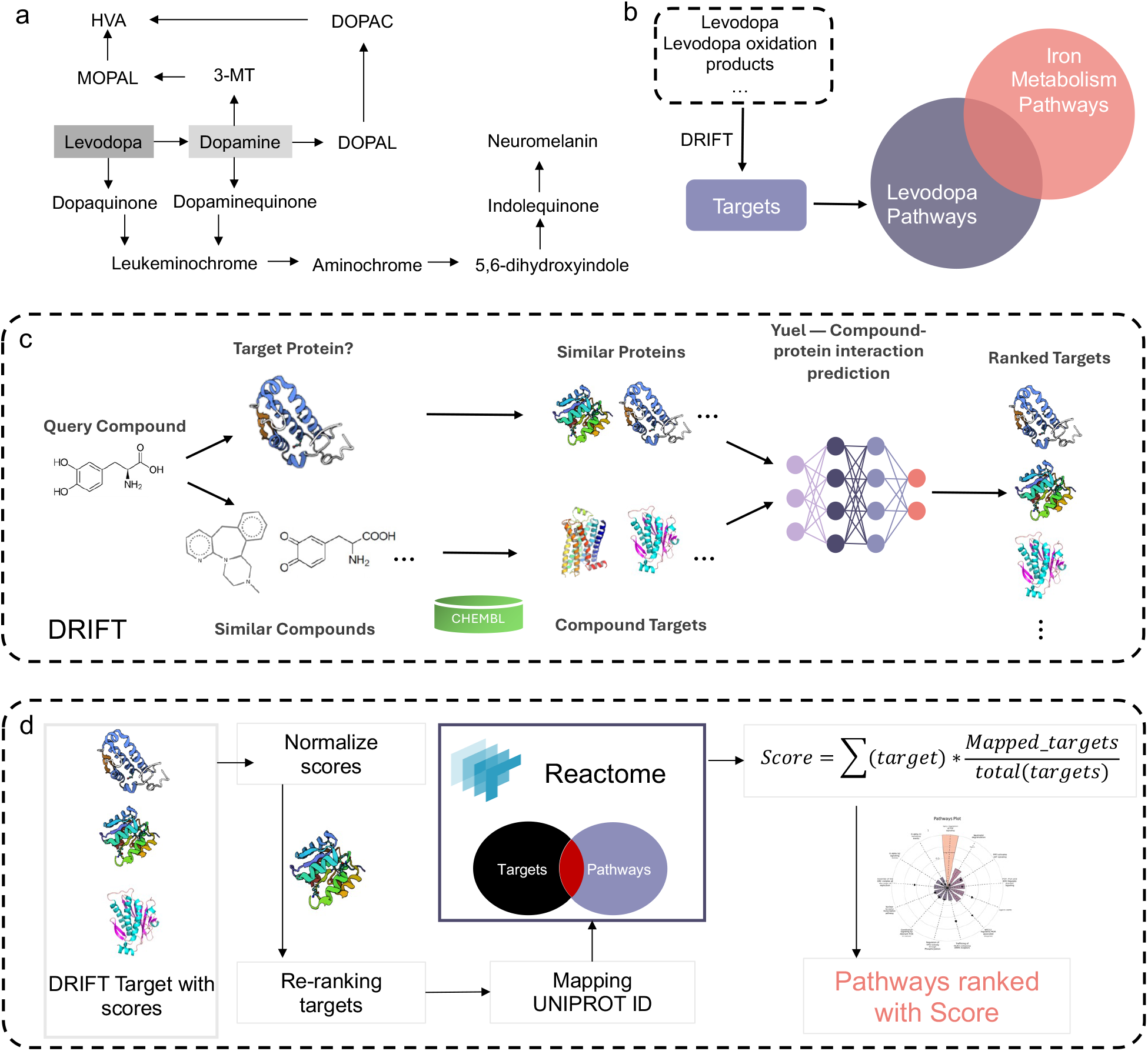
Prediction of the pathways of levodopa and its oxidation products. (a) Levodopa and the oxidation products we tested in our work. The full name of the abbreviations are listed in Table S1. (b) We used DRIFT to predict the targets of levodopa and its oxidation products and then predicted the pathways according to the targets. We identified the targets involved both in the predicted levodopa pathways and the iron metabolism pathways. (c) DRIFT first finds the target proteins of the query compound and then searches for similar proteins of the target proteins. DRIFT also searches for similar compounds of the query compound through databases like CHEMBL. The targets of the similar compounds as well as the similar proteins of the target proteins are subject to Yuel, a compound-protein interaction prediction neural network model. Yuel outputs the ranking of the targets according to their predicted binding affinities to the query compound. (d) The pathways are predicted based on the predicted targets.

**Figure 2.**
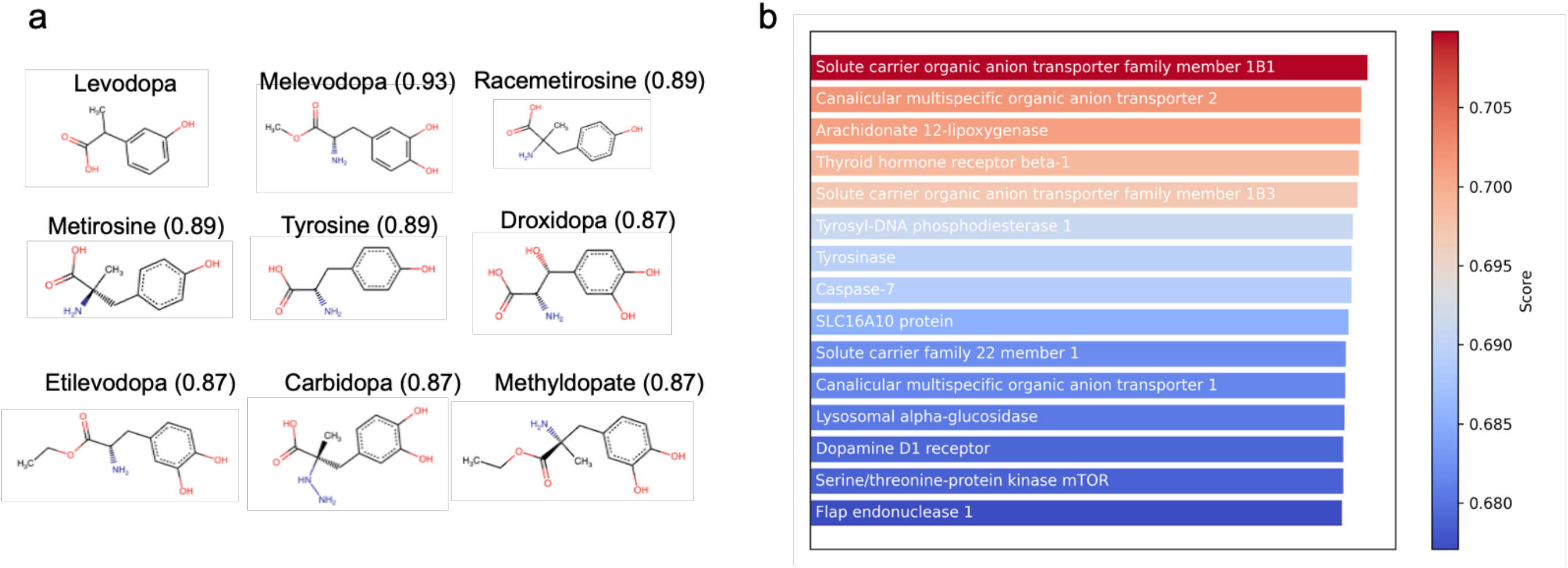
Structurally similar compounds of levodopa and the predicted targets of levodopa. (a) Structurally similar compounds of levodopa obtained via the DRIFT platform by FP2 similarities. (b) The top-15 ranked targets of levodopa predicted by DRIFT.

Since these compounds share key structural elements, they are likely to interact with similar enzymes, transporters, and receptors. For example, some of these compounds are known to interact with aromatic amino acid decarboxylase^25^, OCTs^26^, and solute carriers involved in neurotransmitter transport^27^. Given these similarities, levodopa may engage with the same targets, expanding its potential range of biological interactions.

### Target of levodopa Predicted by DRIFT

The predicted targets of levodopa (**Figure 2B**), including solute carrier transporters (SLCO1B1, SLCO1B3, and SLC22A1), enzymes like tyrosinase, and receptors like the dopamine D_1_ receptor, highlight its broad biological activity. Because the carboxyl group will inhibit isosteric binding to the dopamine D_1_ receptor, it is plausible that levodopa interacts with these receptors via allosteric sites. The interaction of levodopa with solute carriers is especially noteworthy given its structural similarity to related compounds such as droxidopa which also is known to interact with transporters from the OCT family. These solute carriers facilitate the uptake of levodopa and analogs that share similar catecholamine structures. Furthermore, enzymes like tyrosinase and lipoxygenase are involved in metabolic pathways associated with neurotransmitter synthesis and oxidative stress, processes also influenced by structurally similar compounds like metirosine and tyrosine. Tyrosine, for instance, plays a key role in the biosynthesis of catecholamines, and its structural resemblance to levodopa may suggest shared targets in these metabolic pathways. The predicted interaction of levodopa with thyroid hormone receptor beta-1 and mTOR kinase aligns with broader regulatory pathways involved in cell signaling and energy metabolism, areas in which related compounds might also play a role due to their shared functional groups.

The predicted targets of levodopa, such as solute carriers (SLCO1B1, SLCO1B3, SLC22A1) and tyrosinase, may have significant connections to iron metabolism pathways. Tyrosinase, for example, is a copper-dependent enzyme, but it has functional parallels with iron-dependent enzymes involved in melanogenesis, a pathway closely linked to iron homeostasis. Further, disruptions in iron metabolism are often linked to oxidative stress and neurodegenerative processes, areas where enzymes like tyrosinase and lipoxygenase play critical roles. The interaction of levodopa with lipoxygenase, an enzyme that can generate oxidative species, ties levodopa’s action to pathways of cellular oxidative balance, which is directly influenced by iron levels. Moreover, the involvement of solute carriers, especially those in the organic anion transporter families (SLCO1B1, SLCO1B3, and SLC22A1), in transporting a wide variety of substances, including organic ions and iron complexes, suggests that levodopa’s targets may influence iron uptake and distribution. Solute carriers can mediate the transport of iron-bound molecules, and dysfunction in these transporters may lead to altered iron metabolism, affecting cellular processes such as energy production and mitochondrial function. mTOR (mechanistic target of rapamycin), another predicted target, also plays a role in regulating cellular responses to nutrient availability, including iron. Dysregulation of mTOR has been associated with diseases linked to iron overload or deficiency^28^. Furthermore, the thyroid hormone receptor beta-1, which influences metabolic rates, can indirectly affect iron metabolism through its regulation of hemoglobin synthesis and erythropoiesis^29^.

Thus, we performed a detailed analysis of all the iron-related pathways where levodopa is involved. These interactions offer potential mechanisms for how levodopa affects pathways related to iron metabolism.

### Targets of levodopa oxidation products

We predicted the targets of all the levodopa oxidation products listed in Table S1. The predicted targets (**Figure 3**) reveal a broad spectrum of proteins and enzymes, highlighting their diverse roles in cellular processes and their potential connections to iron metabolism. Several of these targets are integral to DNA repair and maintenance, such as Tyrosyl-DNA phosphodiesterase 1, Flap endonuclease 1, DNA polymerase iota, DNA-(apurinic or apyrimidinic site) lyase, and DNA polymerase kappa. These enzymes are crucial for addressing oxidative DNA damage, which is often influenced by iron due to its involvement in generating reactive oxygen species (ROS). The proper functioning of these repair mechanisms is vital to prevent oxidative damage and maintain cellular health, underscoring the link between iron metabolism and cellular repair processes.

**Figure 3.**
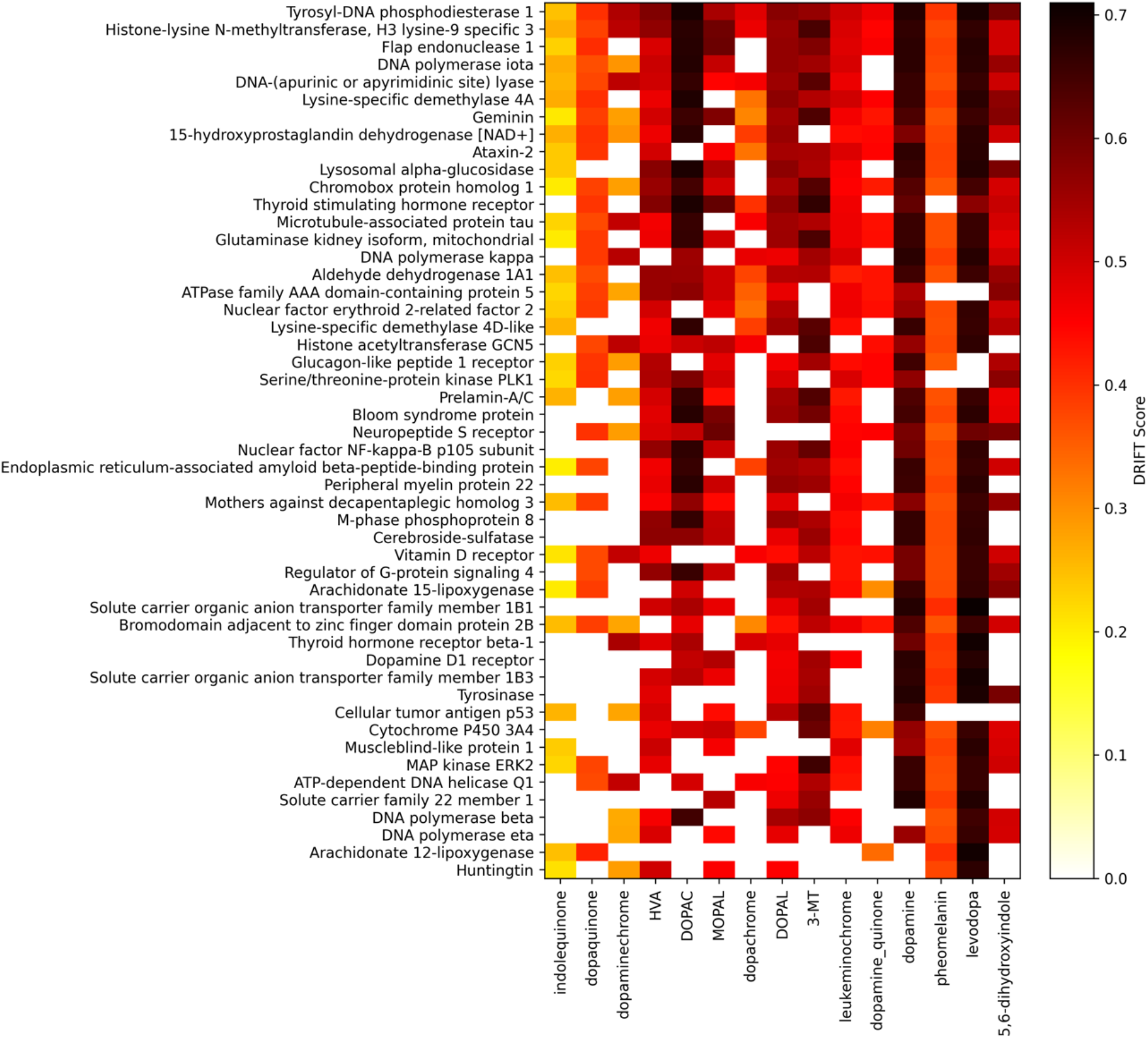
The predicted targets of levodopa and levodopa oxidation products. Only the top 50 ranked are shown.

Transport proteins such as SLCO1B1 and SLC22A1 are crucial for the regulation of nutrient levels, including iron. These transporters help manage the uptake and distribution of iron and other essential nutrients within cells. Their involvement highlights the importance of maintaining proper iron levels and how disturbances in transport mechanisms can impact iron metabolism.

Several of the identified targets are directly involved in metabolic processes that intersect with iron metabolism. For example, 15-hydroxyprostaglandin dehydrogenase [NAD+], Aldehyde dehydrogenase 1A1, and Cytochrome P450 3A4 are important for various metabolic reactions. Cytochrome P450 enzymes, in particular, are involved in the metabolism of heme, an iron-containing molecule, while Aldehyde dehydrogenases help detoxify aldehydes produced through oxidative reactions involving iron. These enzymes’ roles in iron metabolism emphasize the complex interplay between metabolic processes and iron homeostasis.

In addition, transcription factors and cellular response regulators like Nuclear factor erythroid 2-related factor 2 (Nrf2) and Nuclear factor NF-kappa-B p105 subunit play pivotal roles in managing cellular responses to stress and inflammation, processes that are closely linked to iron metabolism. Nrf2, in particular, regulates genes that respond to oxidative stress and can affect iron-related gene expression. Similarly, NF-kappa-B is involved in inflammatory responses, which can intersect with iron metabolism pathways.

### Pathway Analysis

We utilized a pathway analysis algorithm (Figure 4; Methods) to predict the pathways of levodopa and its metabolites according to their predicted targets (Figure S1-S14). The algorithm revealed several pathways intricately linked to iron metabolism.

**Figure 4.**
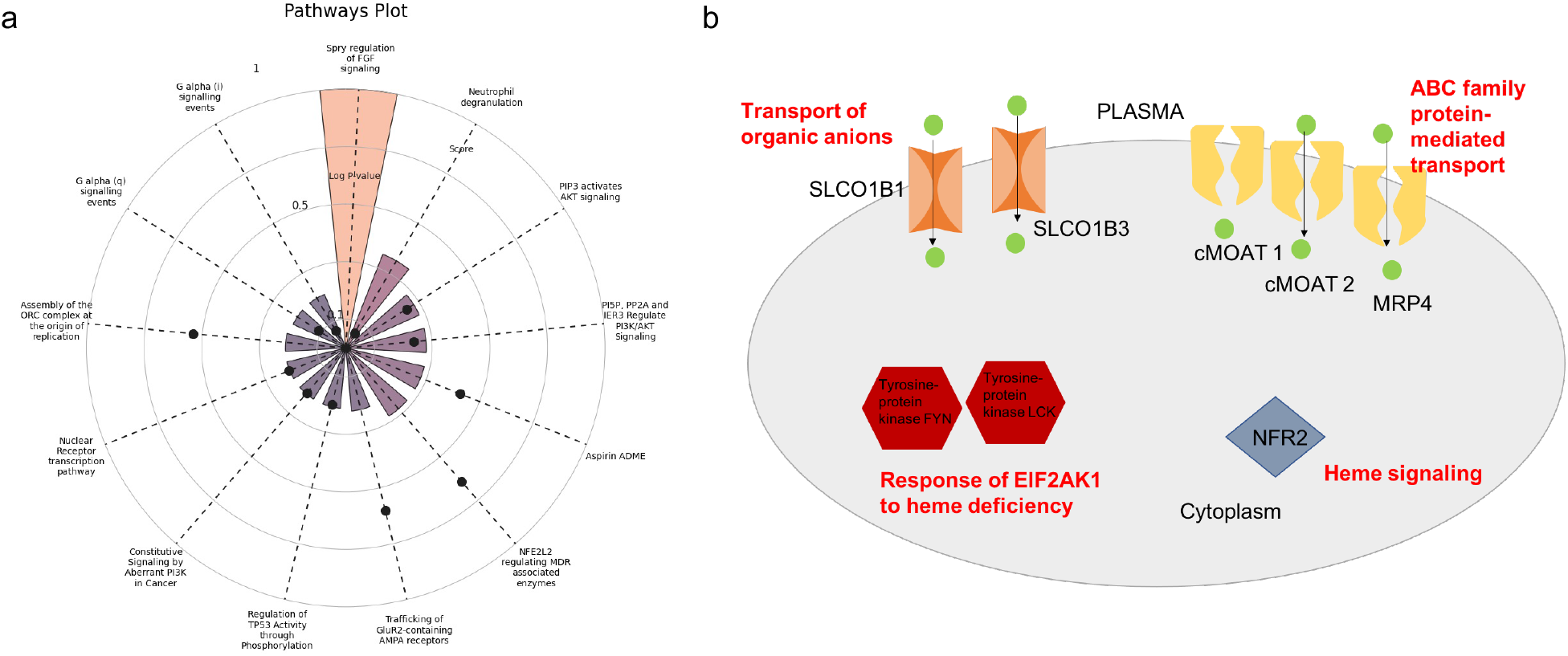
Pathways analysis of levodopa. (a) The top-ranked pathways of levodopa predicted by the pathway analysis algorithm. (b) The levodopa pathways are related to iron metabolism. Predicted targets of levodopa are labeled with black color. Pathways are labeled by red color.

#### Heme signaling pathway

Nuclear factor erythroid 2-related factor 2 (NRF2) has emerged as a critical protein connected to the heme signaling pathway, highlighting its importance in iron regulation^30^. Recent studies on honey and levodopa have provided insights into their potential neuroprotective effects through modulation of the NRF2 signaling pathway. A study comparing honey and levodopa in an MPTP-induced PD model demonstrated that both substances protected substantia nigra pars compacta (SNpc) neurons against oxidative stress by modulating the ROS-Nrf2-GSH pathway. This finding suggests that honey and levodopa can upregulate antioxidant markers, including glutathione (GSH) and NRF2, potentially protecting against oxidative stress-induced neuronal damage in PD. Additionally, levodopa has been found to form stable complexes with iron, potentially altering iron deposition and transport. These findings collectively highlight the intricate relationships between heme signaling, NRF2 activation, iron homeostasis, and dopamine metabolism in the context of PD.

#### Response of EIF2AK1 to heme deficiency pathway

The heme-regulated inhibitor (HRI) kinase is activated in response to heme deficiency and is directly linked to iron metabolism, as heme is an iron-containing molecule^31^. The predicted interaction of levodopa with kinases (e.g., tyrosine-protein kinase FYN and tyrosine-protein kinase LCK) may influence the HRI pathway, potentially altering the cellular response to iron deficiency. By modulating this pathway, levodopa could help cells adapt to iron scarcity, possibly leading to changes in iron uptake or utilization to maintain cellular functions.

#### ABC-family proteins mediated transport pathway

ABC transporters, including canalicular multispecific organic anion transporter 1 and canalicular multispecific organic anion transporter 2, transport a variety of substances, including metals like iron^32^. If levodopa modulates these transporters, it could affect the movement of iron-containing organic molecules, influencing both iron uptake and export. This interaction may play a role in maintaining iron balance, especially in tissues where iron transport is crucial for metabolic processes.

#### Transport of organic anions/cations pathway

Solute carrier proteins, such as SLCO1B1 and SLCO1B3, are known to transport organic anions, including iron bound to organic molecules^33^. By interacting with these transporters, levodopa may influence the distribution of iron within the body, potentially affecting iron availability at the cellular level. This could have significant implications for tissues that require precise regulation of iron, such as the brain and liver, where iron homeostasis is critical for neurotransmitter synthesis and energy production. Further, SLC22A1, one of the predicted targets of levodopa, plays a significant role in the transport of organic cations and is closely related to iron metabolism. SLC22A1 shares structural similarities with SLC22A17, which is the receptor for lipocalin-2 (LCN2), a protein involved in iron transport and regulation^34^. Lipocalin-2 sequesters iron by binding to siderophores, and its interaction with SLC22A17 regulates iron uptake and release in various tissues, particularly during inflammation and cellular stress. This connection suggests that SLC22A1 may also influence iron homeostasis indirectly, possibly through similar mechanisms of organic cation transport that overlap with iron metabolism pathways.

Through these pathways, levodopa may influence iron metabolism, impacting how cells manage iron availability, storage, and detoxification. This could be particularly relevant in neurodegenerative conditions like Parkinson’s disease, where both iron dysregulation and oxidative stress are contributing factors.

## Discussion

It was recently reported that levodopa had an affinity for siderocalin (Scn)/lipocalin-2 by forming a stable complex in the presence of iron.^35^ This complex may facilitate cellular iron uptake, potentially leading to iron accumulation in cells that then would increase oxidative stress and downstream effects like neuroinflammation. These data supported the hypothesis that while levodopa has essential benefits on brain health by normalizing circuitry, it causes concomitant cellular damage. In fact, this idea reconciles two apparently contradictory findings: that levodopa is essential for patient welfare and brain health ^10,36^, but causes decreases in striatal dopamine terminal density suggestive of neuronal damage^37^.

The findings of this study provide new insights into the molecular interactions of levodopa and its metabolites, particularly its potential role in modulating iron metabolism in Parkinson’s disease (PD). Levodopa has long been recognized as a double-edged sword in PD treatment. While it is the most effective drug for managing motor symptoms, its prolonged use is associated with side effects that may accelerate neurodegeneration. Our study sheds light on the potential mechanisms underlying these adverse effects, specifically through interactions with proteins involved in iron homeostasis. One of the most intriguing results from our analysis is the possible interaction between levodopa and SLC22A17 (24p3R) (**Figure 5**), a receptor involved in iron uptake through its binding with lipocalin-2. Although DRIFT did not predict this interaction directly, the structural similarity between SLC22A1, a predicted target of levodopa, and SLC22A17 raises the possibility that levodopa could interact with SLC22A17 as well. If levodopa interacts with SLC22A17, it could enhance cellular iron uptake, thereby contributing to the observed iron accumulation in PD patients. This hypothesis is particularly compelling in existing literature that links increased brain iron levels with PD progression.

**Figure 5.**
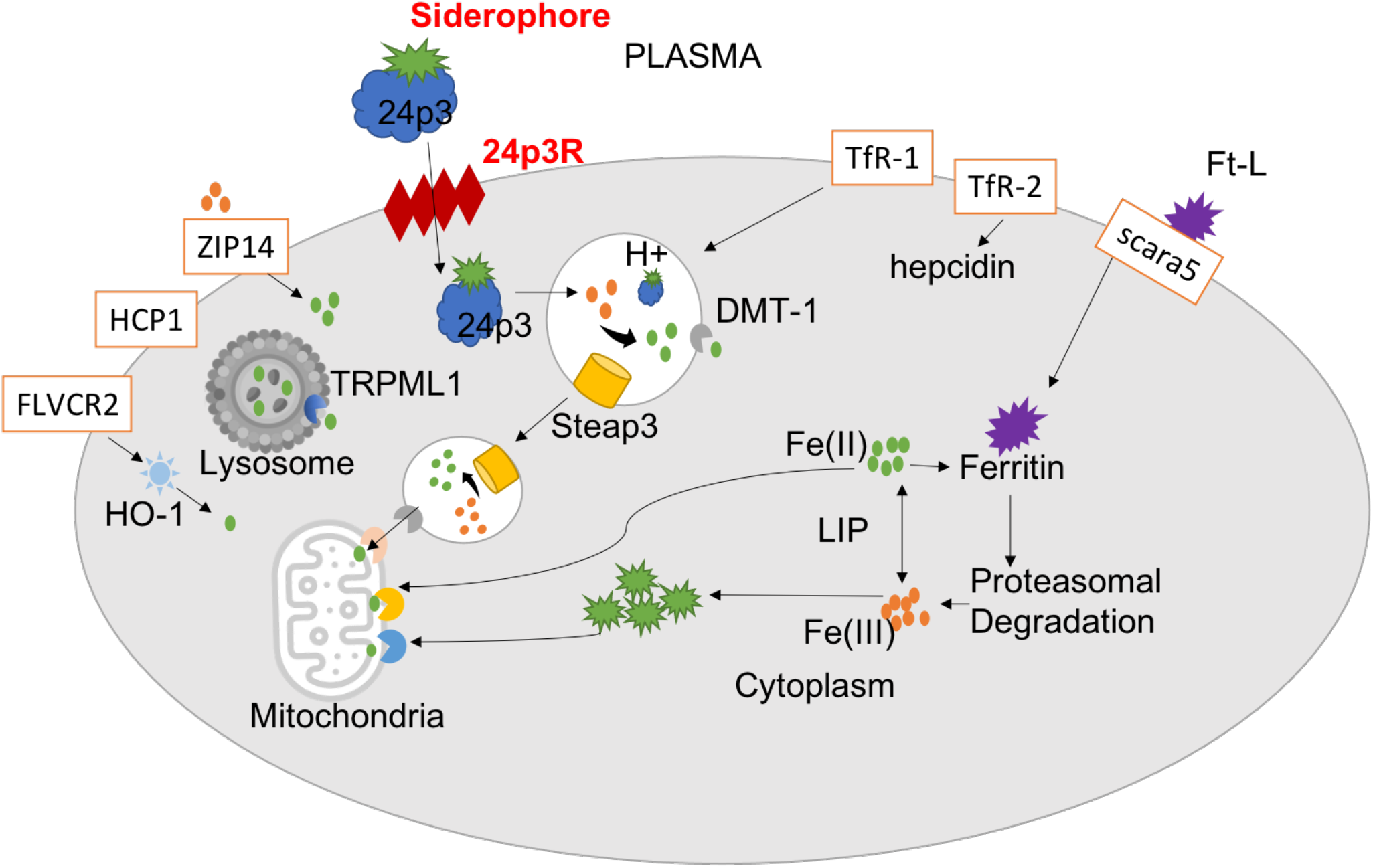
The iron metabolism pathways. Levodopa is predicted to interact with SLC22A17, which may enhance cellular iron uptake by binding 24p3/siderophore complex, thereby contributing to iron uptake.

The hypothesis that levodopa interacts with iron-related pathways opens up several new avenues for research. If confirmed, these interactions could lead to a reevaluation of current PD treatment strategies, particularly in the context of long-term levodopa use. For instance, developing adjunct therapies that mitigate levodopa’s impact on iron metabolism could enhance the drug’s safety profile and reduce the risk of neurodegeneration. Moreover, our findings underscore the importance of considering off-target effects when evaluating the safety and efficacy of medications like levodopa. While levodopa’s primary mode of action is through its conversion to dopamine, its interactions with other proteins and pathways could have significant, albeit unintended, consequences. Understanding these off-target effects will be crucial for developing more effective and safer therapies for PD.

## Conclusion

In summary, our study suggests that levodopa may interact with proteins and pathways involved in iron metabolism, potentially contributing to the neurotoxic side effects observed in PD patients. These findings highlight the need for a deeper understanding of the molecular mechanisms of levodopa and pave the way for future research aimed at improving the safety and efficacy of PD treatments.

## Methods

### DRIFT (Drug Target Identification)

DRIFT is a computational pipeline designed to identify protein binders for any active chemical molecule on a proteome-wide scale^24^. DRIFT leverages both two-dimensional (2D) and three-dimensional (3D) chemical similarity searches to find compounds that are structurally similar to a query molecule. It then ranks the associated protein targets using an attention-based neural network. This process involves the generation of pharmacophore and molecular fingerprints for the query compound, which are then compared to a curated database of annotated compounds using Tanimoto coefficients for similarity scoring. The top-ranked targets are subsequently analyzed for their biological functions and processes through Gene Ontology enrichment. DRIFT’s web server provides a user-friendly interface for researchers to submit queries, view results, and explore chemically similar compounds and their potential targets.

### Target Prediction Using DRIFT

The chemical structure of levodopa was provided as input to DRIFT. The platform utilizes both 2D and 3D chemical descriptors to capture the molecular features relevant to binding interactions. Additionally, known protein targets of structurally similar compounds were included as reference data to guide the prediction process. DRIFT uses a combination of cheminformatics and machine learning methods to map the interaction landscape of levodopa across the proteome. The platform predicts potential protein targets based on binding affinity scores, which are calculated through Yuel, which is an attention-based neural network model. The results are ranked according to the likelihood of interaction, with special emphasis on proteins related to iron metabolism.

### Pathway Prediction

Details of the pathway analysis algorithm are described in reference^38^. Here, we give a brief introduction to the algorithm. Initially, the DRIFT-generated target scores are normalized and used as input for pathway elucidation. We utilize the Reactome database, a comprehensive, freely available repository of biological pathways, to map the identified targets to relevant pathways and rank them based on their relevance to the targets. The output presents a list of pathways, along with their associated targets and corresponding scores, providing a comprehensive view of the compound-to-pathway relationships.

## Acknowledgments

The author(s) declare that financial support was received for the research, authorship, and/or publication of this article. We acknowledge support from the National Institutes of Health (NIH) 1R35 GM134864 (ND), 1R01 AG071675 (ND, RM), and the Passan Foundation ND).

