## Supplementary Information for "Comprehensive mapping of the Interaction of levodopa and iron metabolism in Parkinson’s disease"

**Table S1. Levodopa and levodopa metabolites tested by DRIFT.**

| **Compound Name** | **Abbreviation** | **SMILES** |
| --- | --- | --- |
| Dopamine | DA | C1=CC(=C(C=C1CCN)O)O |
| Levodopa | DOPA | C1=CC(=C(C=C1C[C@@H](C(=O)O)N)O)O |
| 3-methoxy-4-hydroxyphenylacetaldehyde | MOPAL | COC1=C(C=CC(=C1)C(C=O)O)O |
| 3,4-dihydroxyphenylacetic acid | DOPAC | C1=CC(=C(C=C1CC(=O)O)O)O |
| 3,4-dihydroxyphenylacetaldehyde | DOPAL | C1=CC(=C(C=C1CC=O)O)O |
| dopaquinone |  | C1=CC(=O)C(=O)C=C1C[C@@H](C(=O)O)N |
| dopamine quinone |  | C1=CC(=O)C(=O)C=C1CCN |
| Indolequinone |  | C1=CNC2=CC(=O)C(=O)C=C21 |
| Leukeminochrome |  | C1CNC2=CC(=C(C=C21)O)O |
| Dopachrome |  | C1C(NC2=CC(=O)C(=O)C=C21)C(=O)O |
| Dopaminechrome/Aminochrome |  | C1CNC2=CC(=O)C(=O)C=C21 |
| 5,6-dihydroxyindole |  | C1=CNC2=CC(=C(C=C21)O)O |
| Homovanillic acid | HVA | COC1=C(C=CC(=C1)CC=O)O |
| 3-methoxytyramine | 3-MT | COC1=C(C=CC(=C1)CCN)O |
| Pheomelanin monomer |  | NC(CSC1=CC(CC(N)C(O)=O)=CC(O)=C1O)C(O)=O |

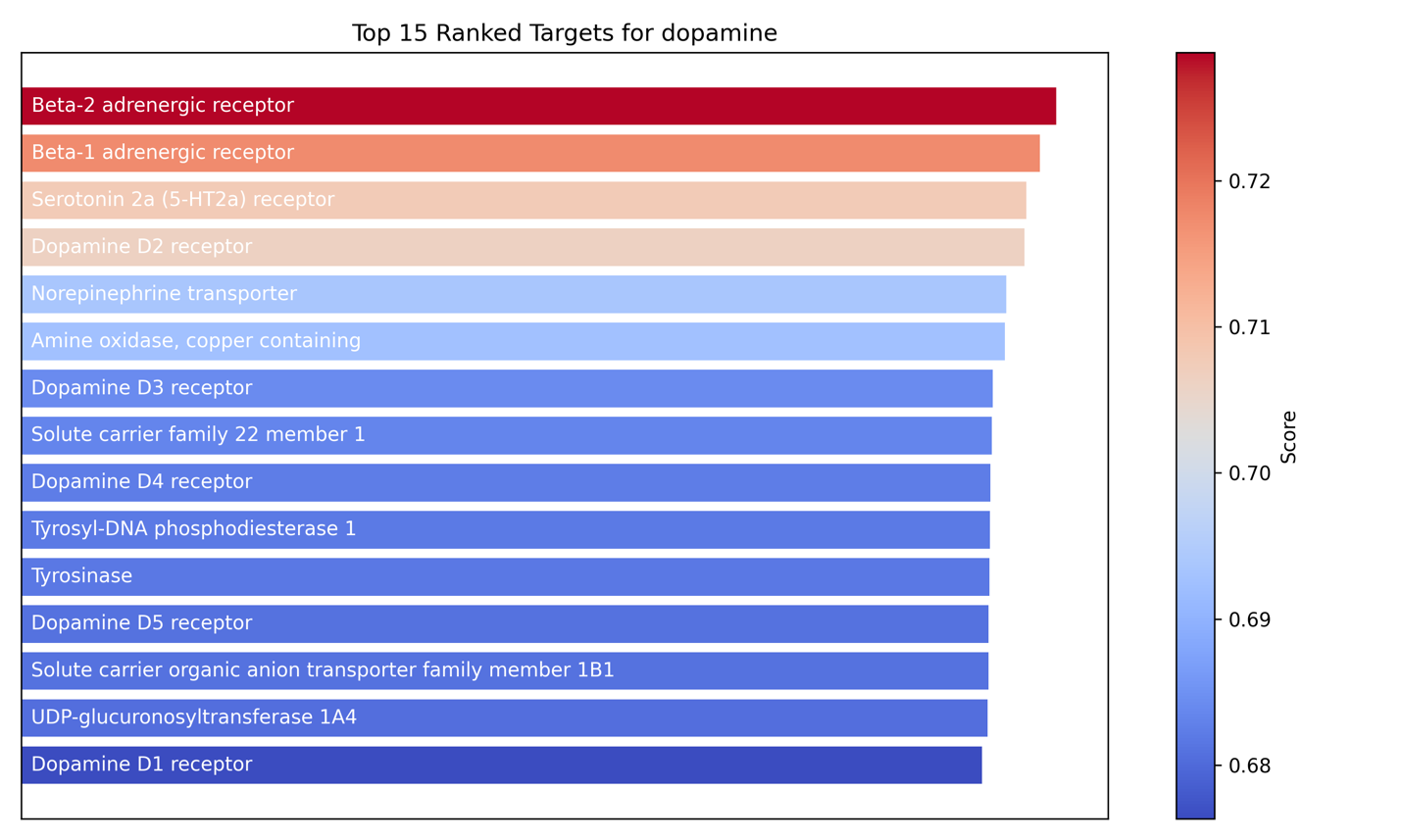

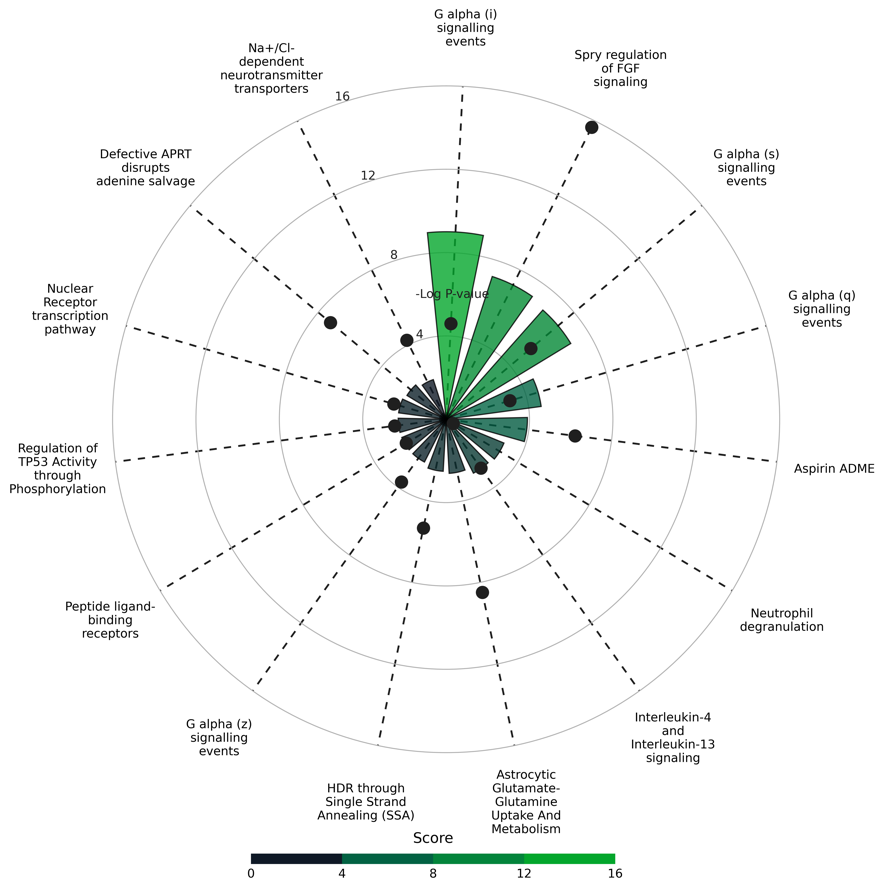

Figure S1. **Top 15 ranked targets predicted and mapped pathways for dopamine.**

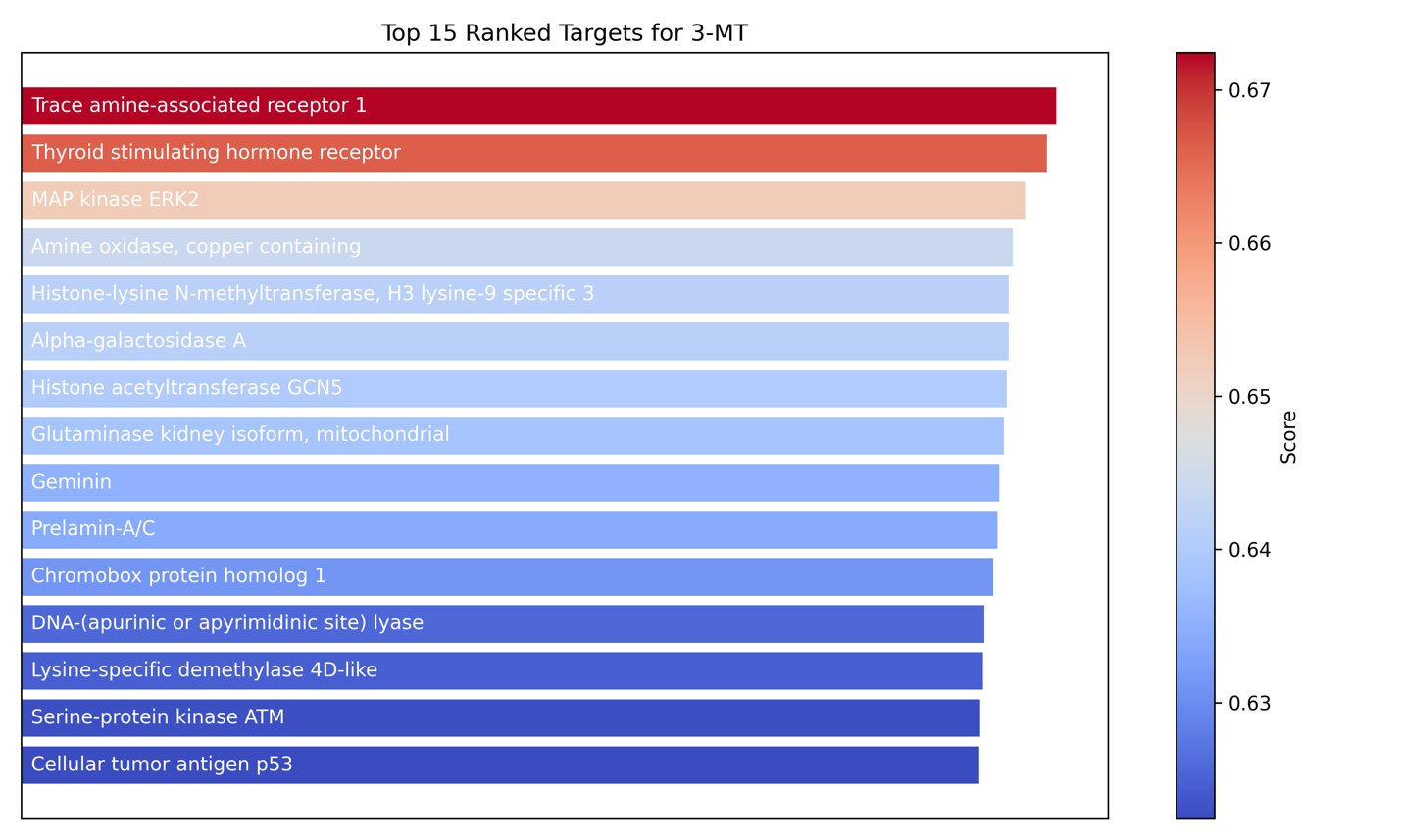

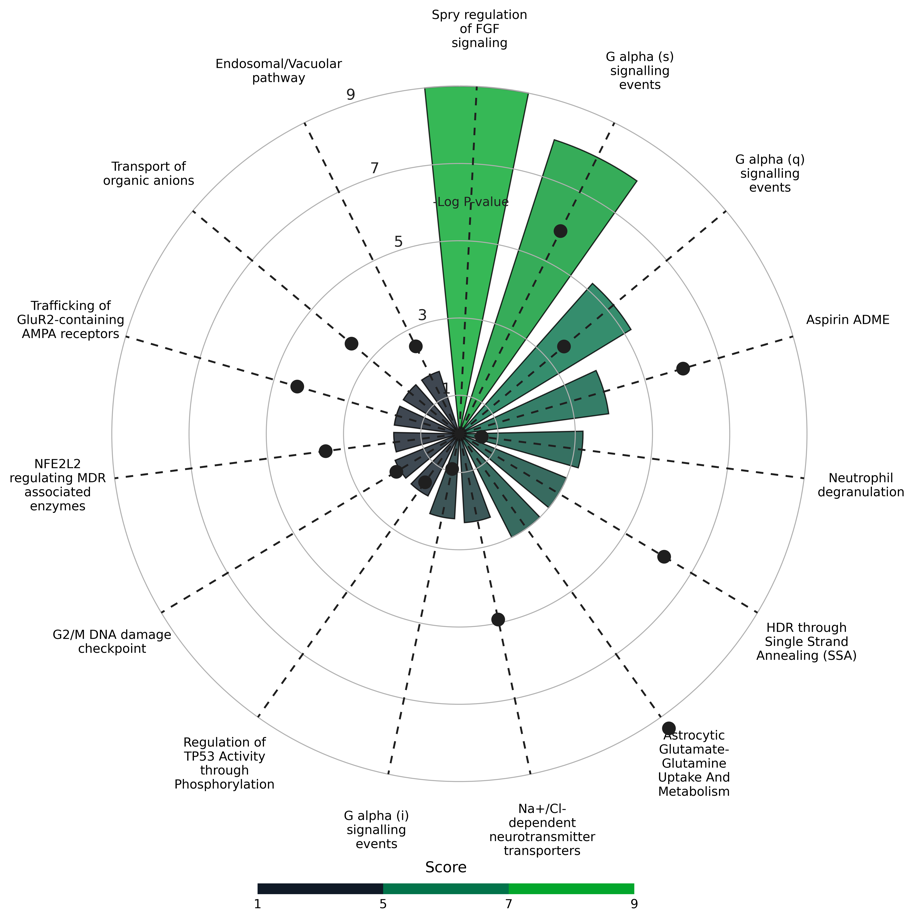

Figure S2. **Top 15 ranked targets predicted and mapped pathways for 3-MT.**

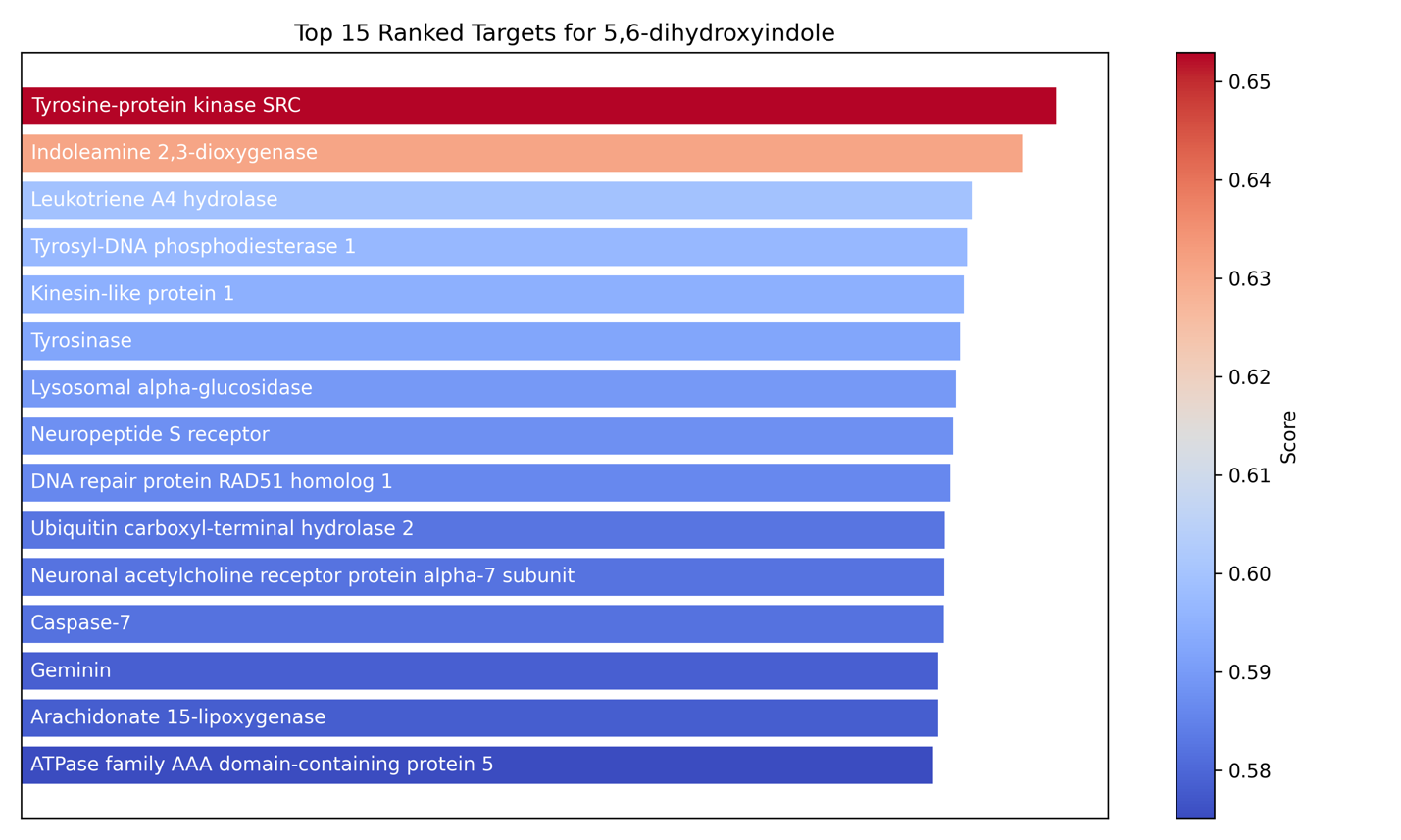

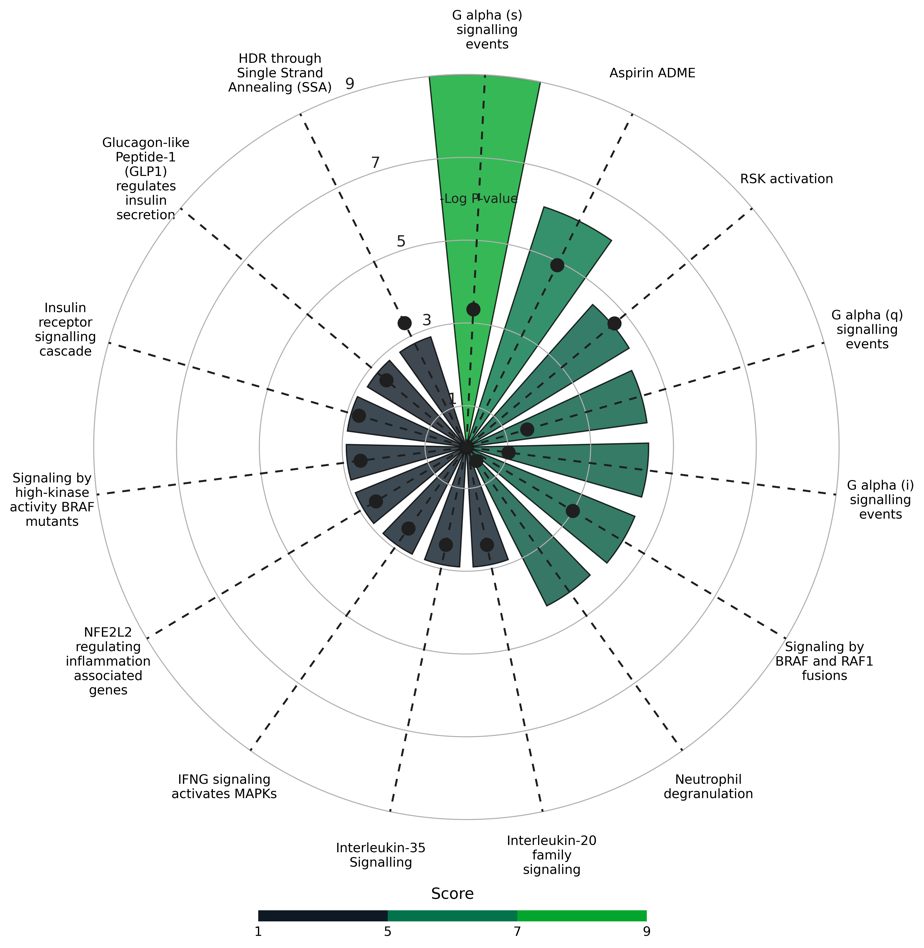

Figure S3. **Top 15 ranked targets predicted and mapped pathways for 5,6-dihydroxyindole.**

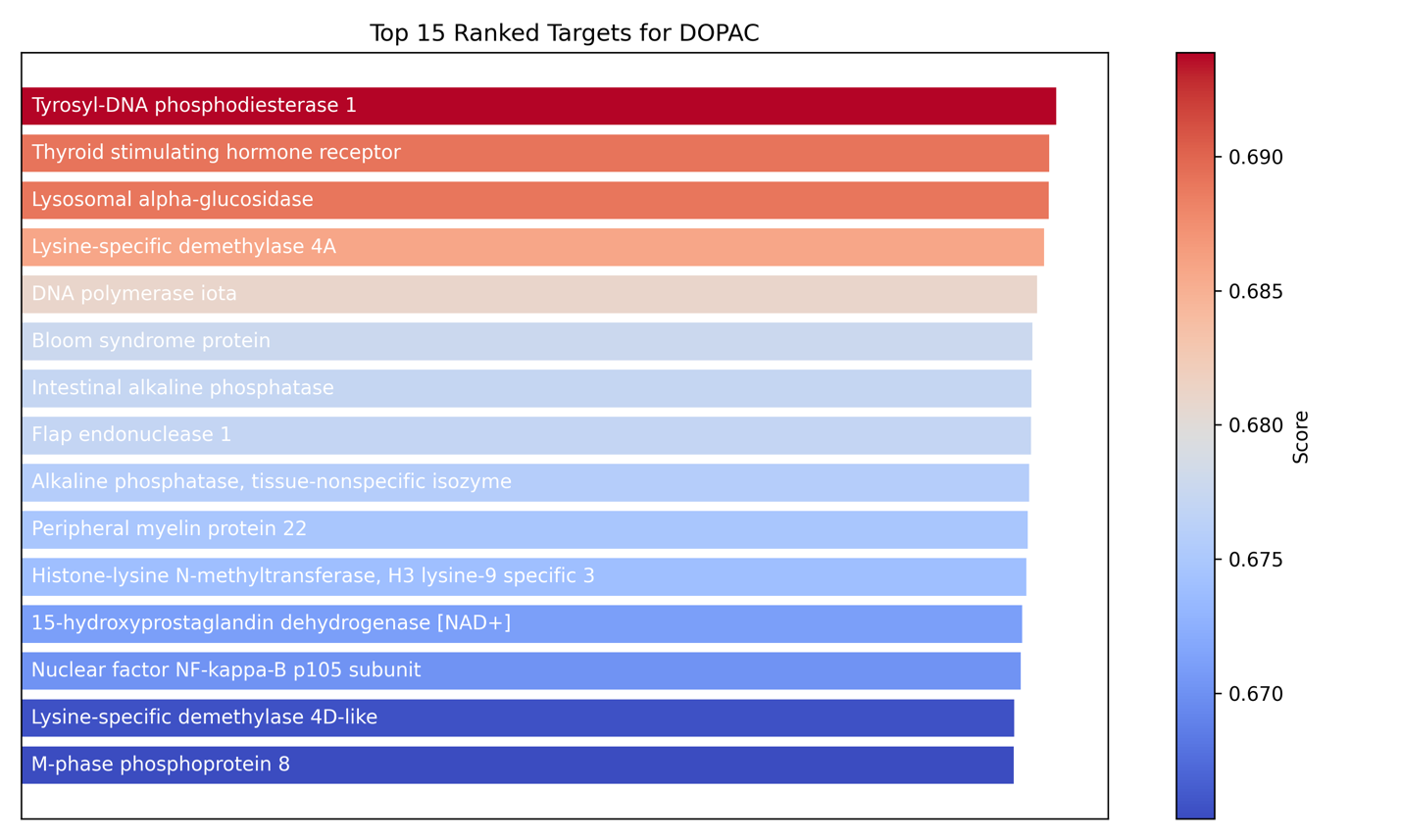

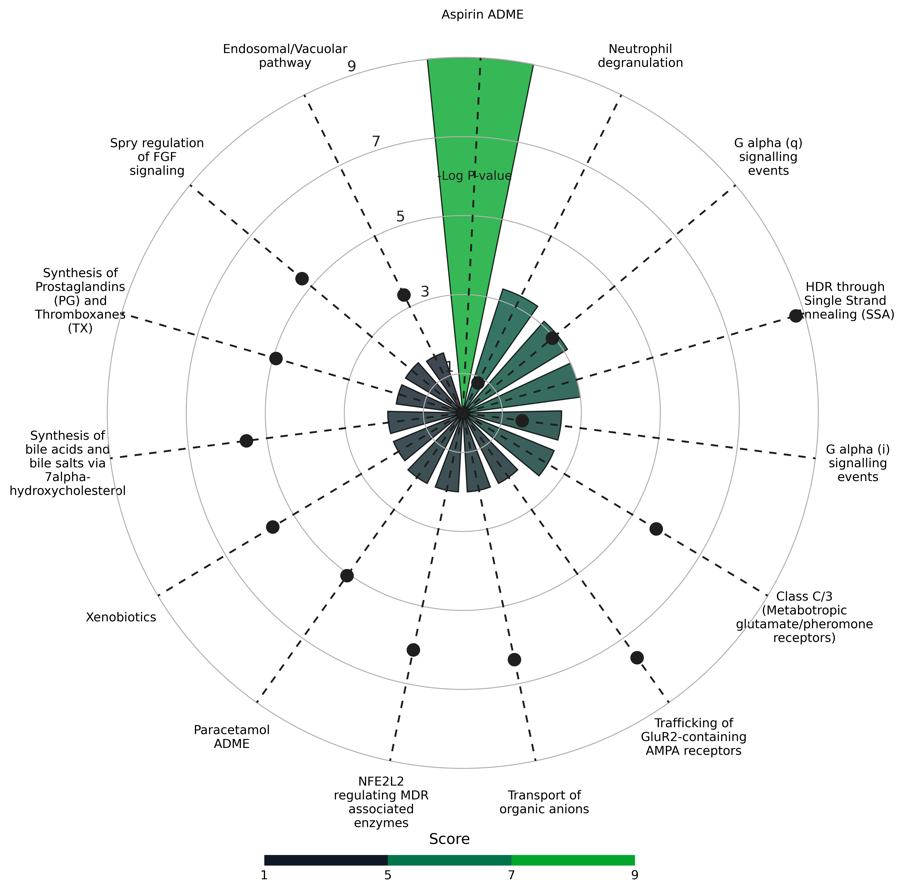

Figure S4. **Top 15 ranked targets predicted and mapped pathways for DOPAC.**

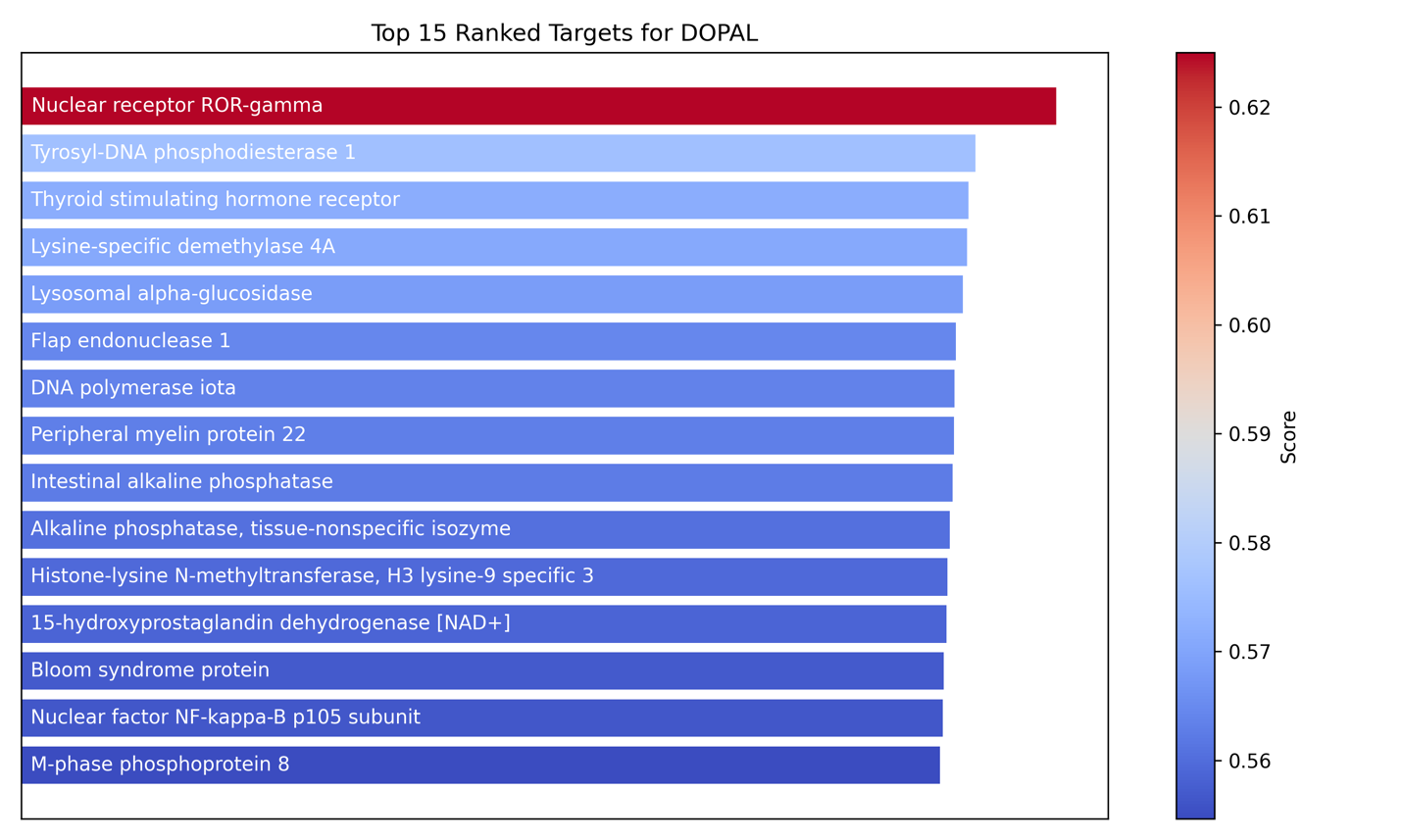

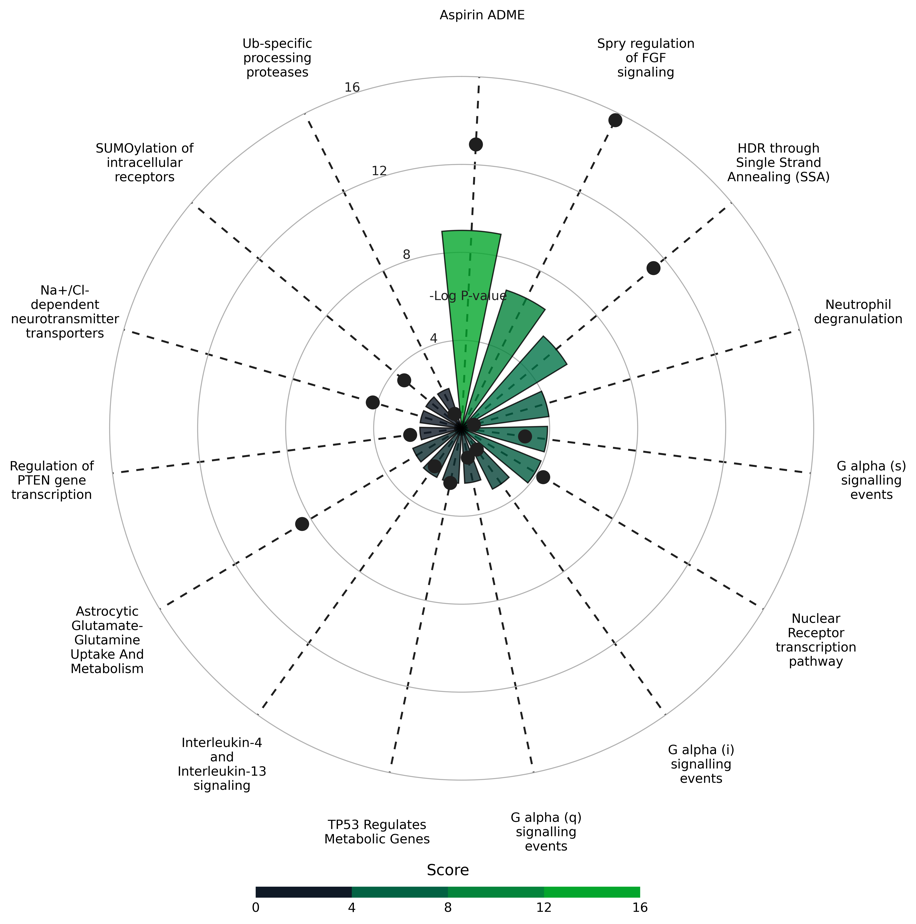

Figure S5. **Top 15 ranked targets predicted and mapped pathways for DOPAL.**

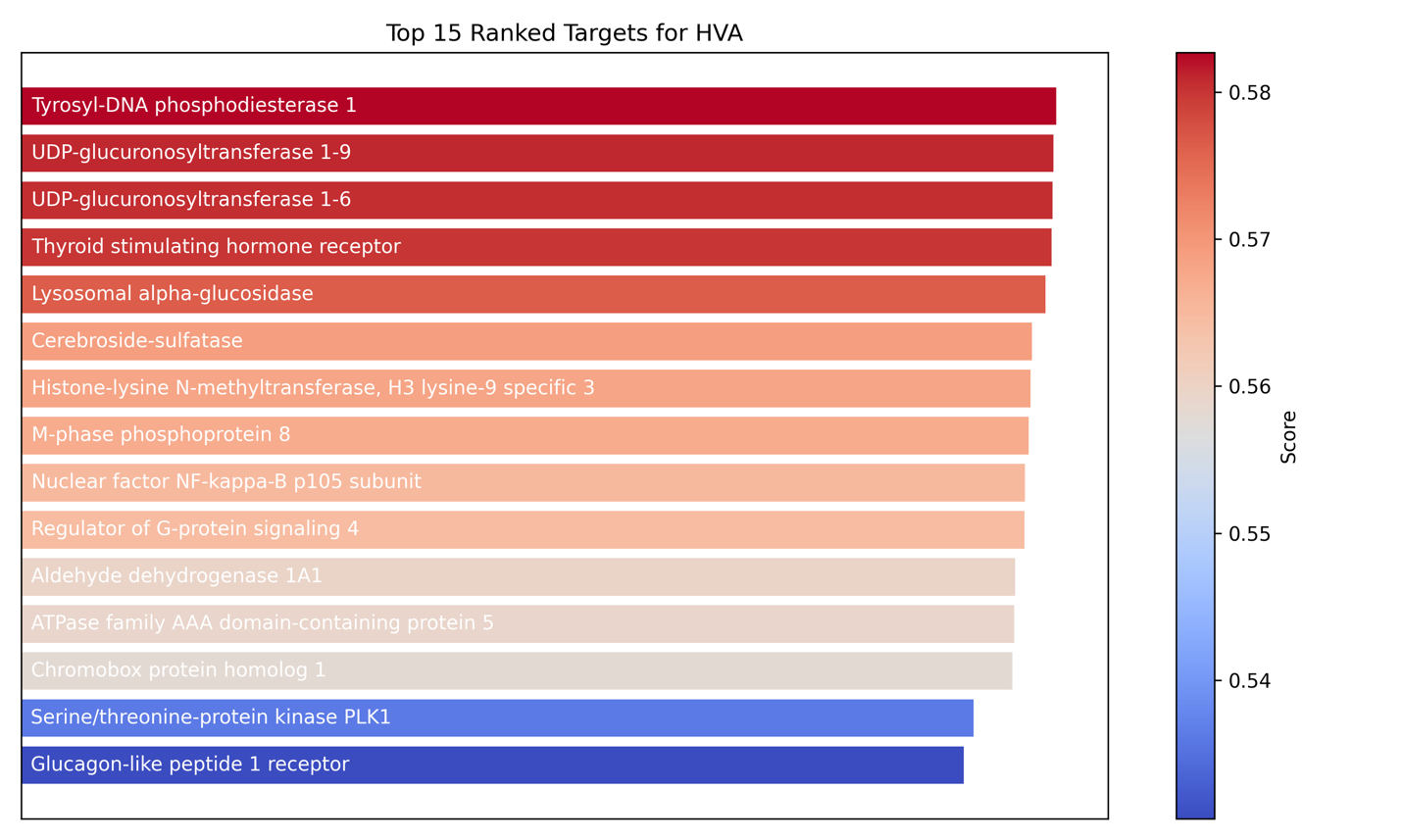

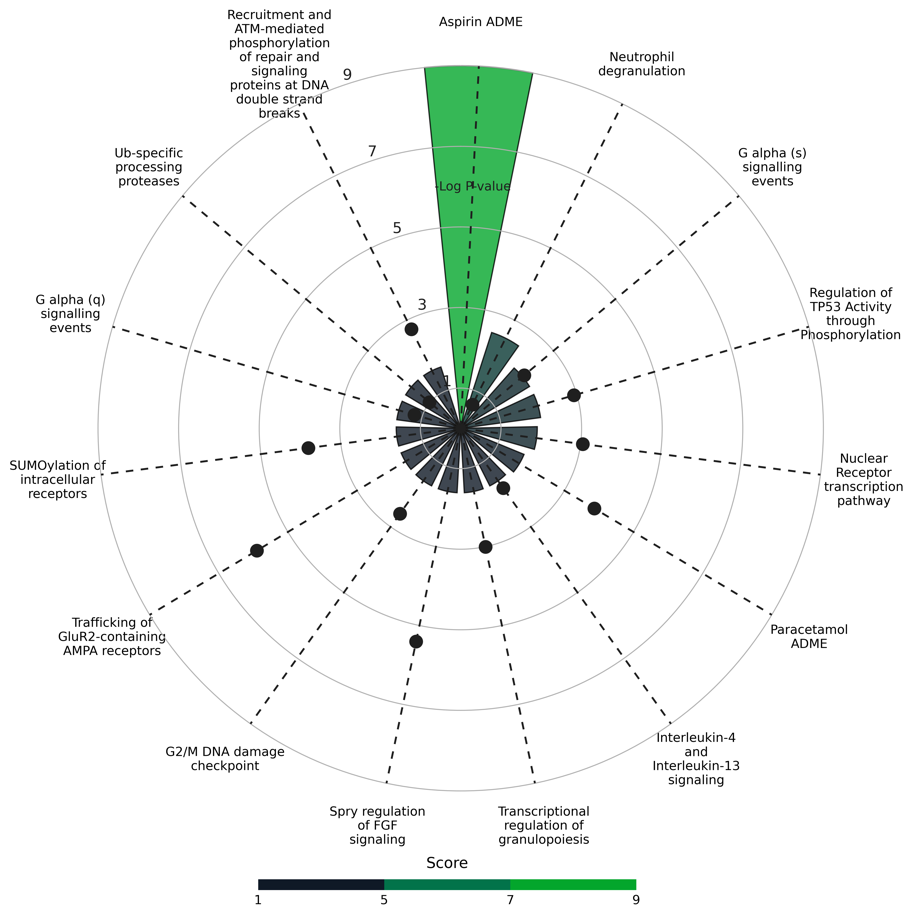

Figure S6. **Top 15 ranked targets predicted and mapped pathways for HVA.**

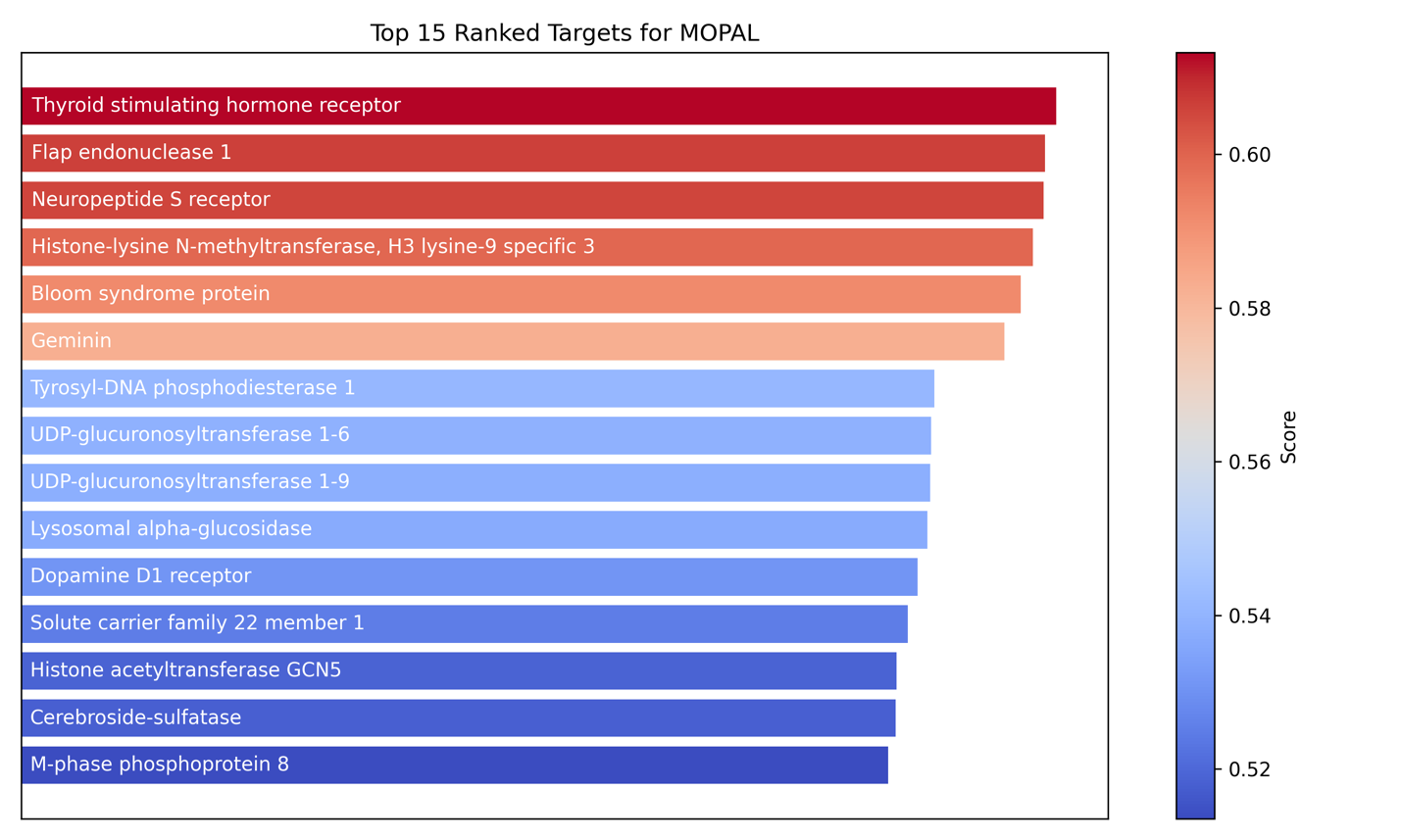

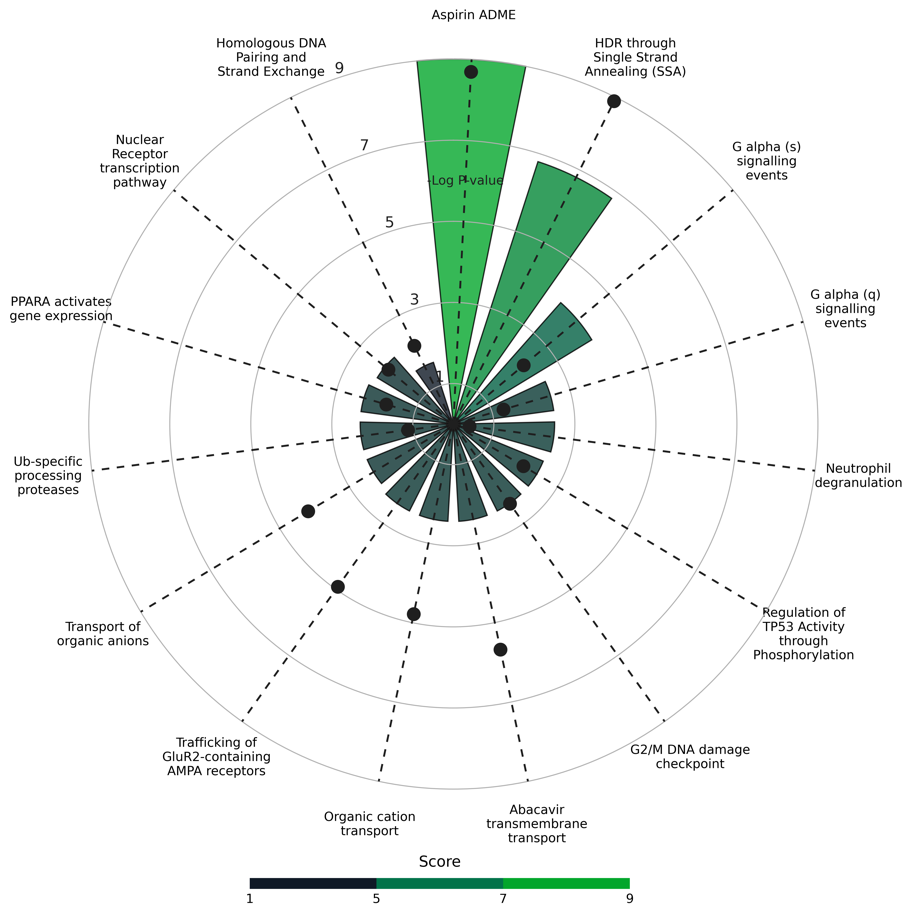

Figure S7. **Top 15 ranked targets predicted and mapped pathways for MOPAL.**

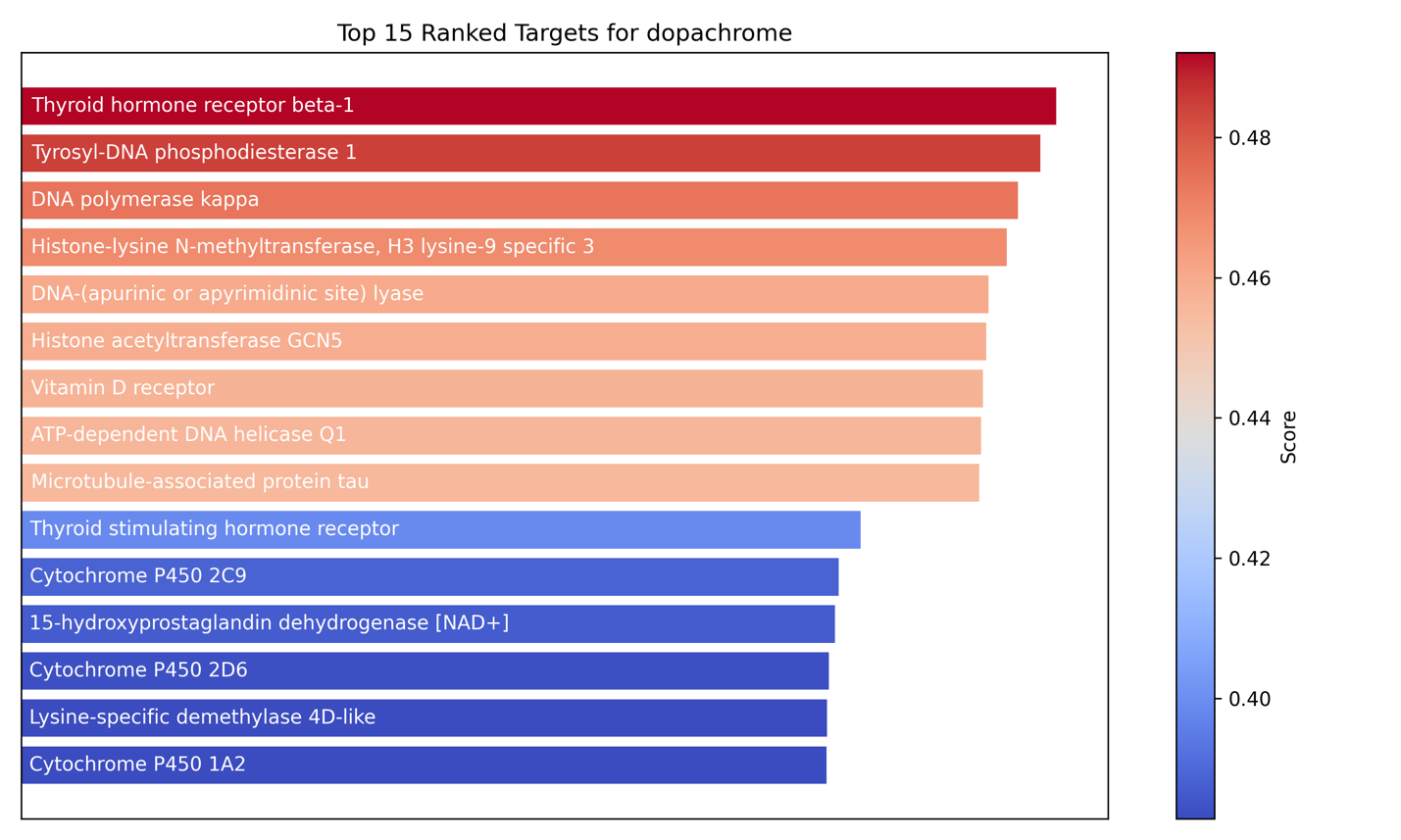

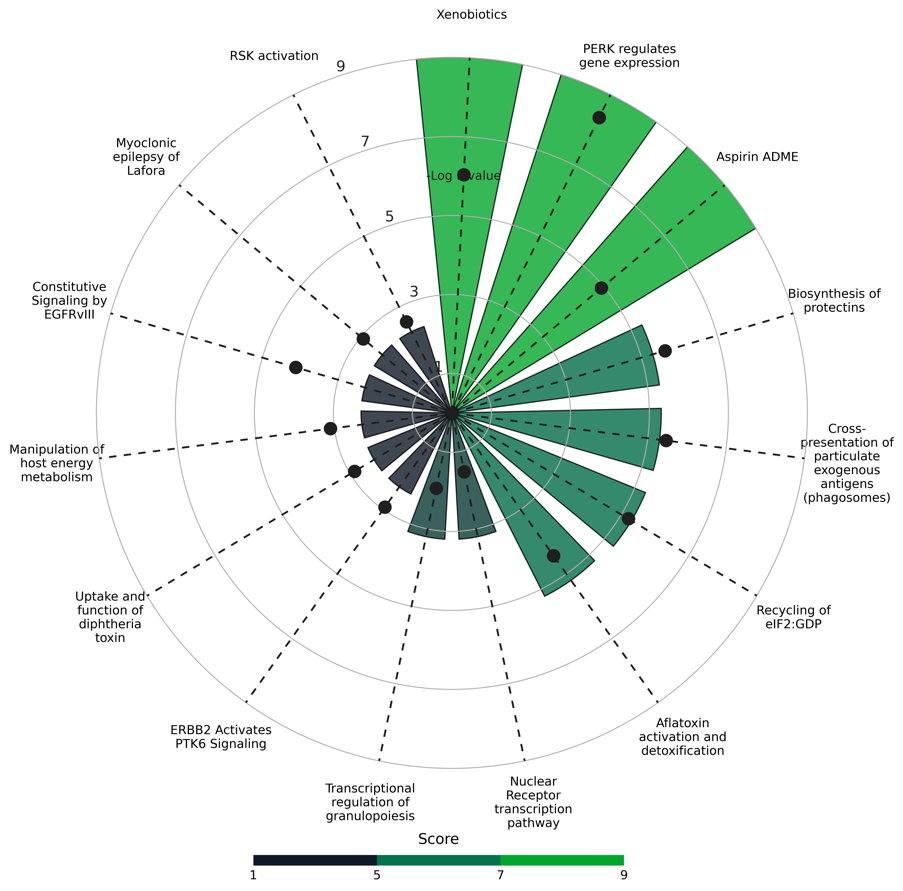

Figure S8. **Top 15 ranked targets predicted and mapped pathways for dopachrome.**

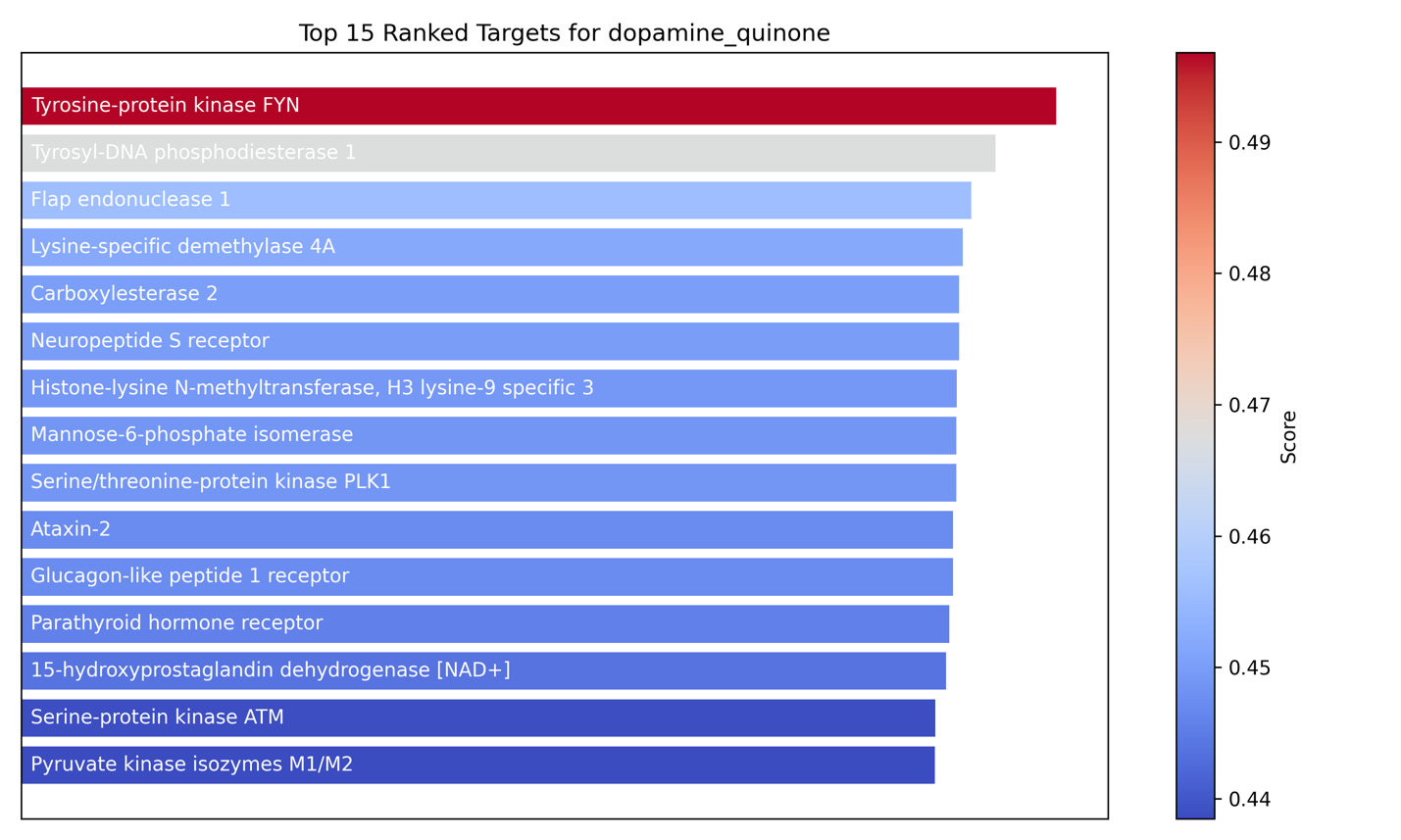

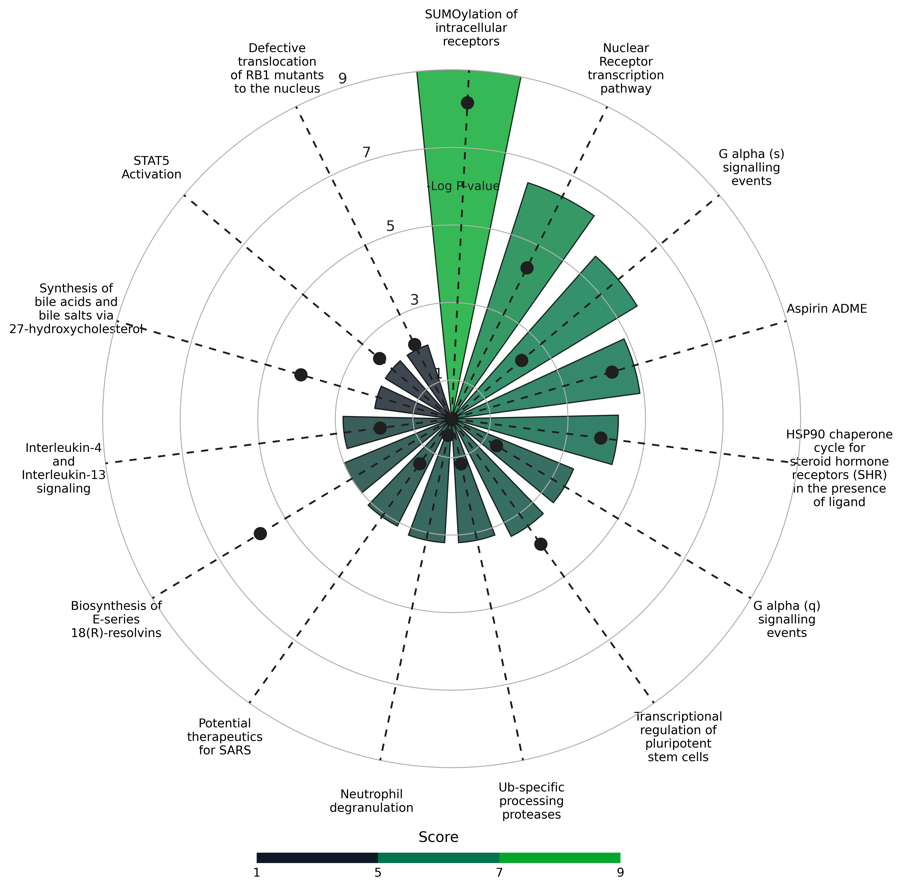

Figure S9. **Top 15 ranked targets predicted and mapped pathways for dopamine quinone.**

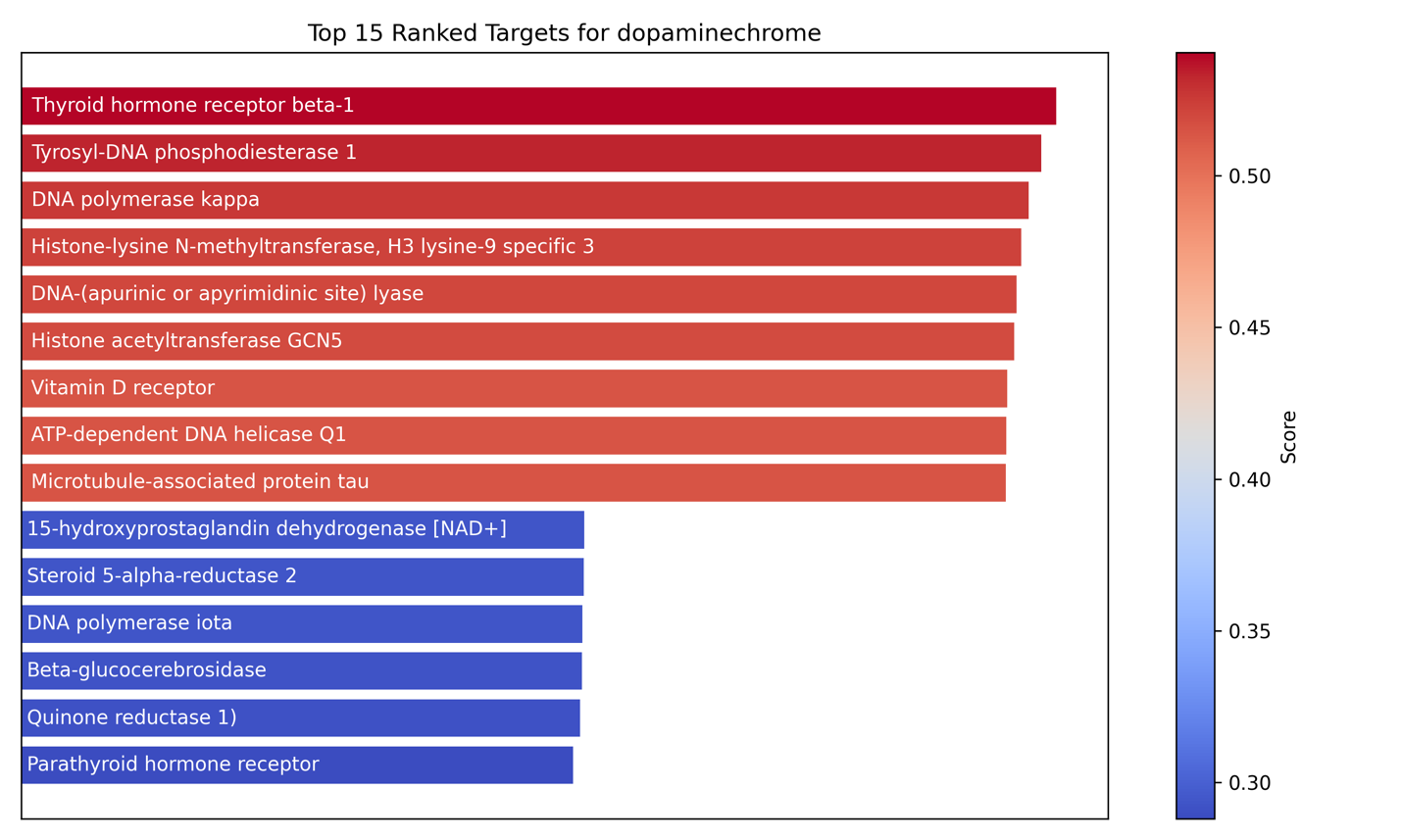

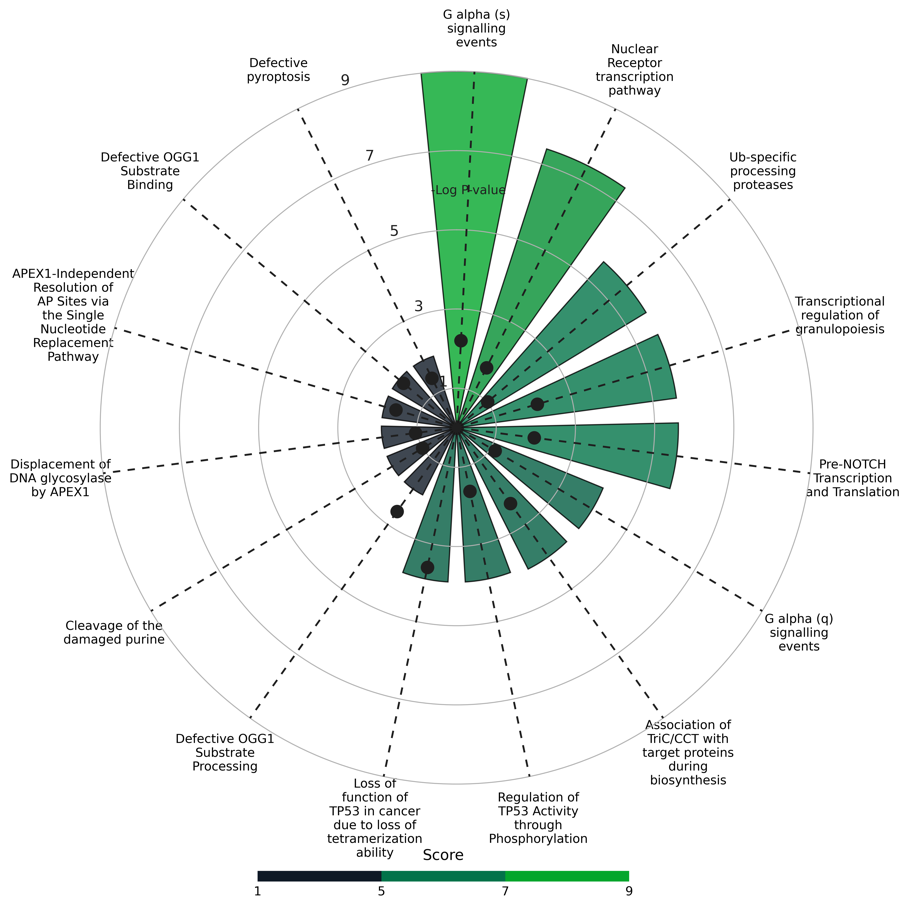

Figure S10. **Top 15 ranked targets predicted and mapped pathways for dopaminechrome.**

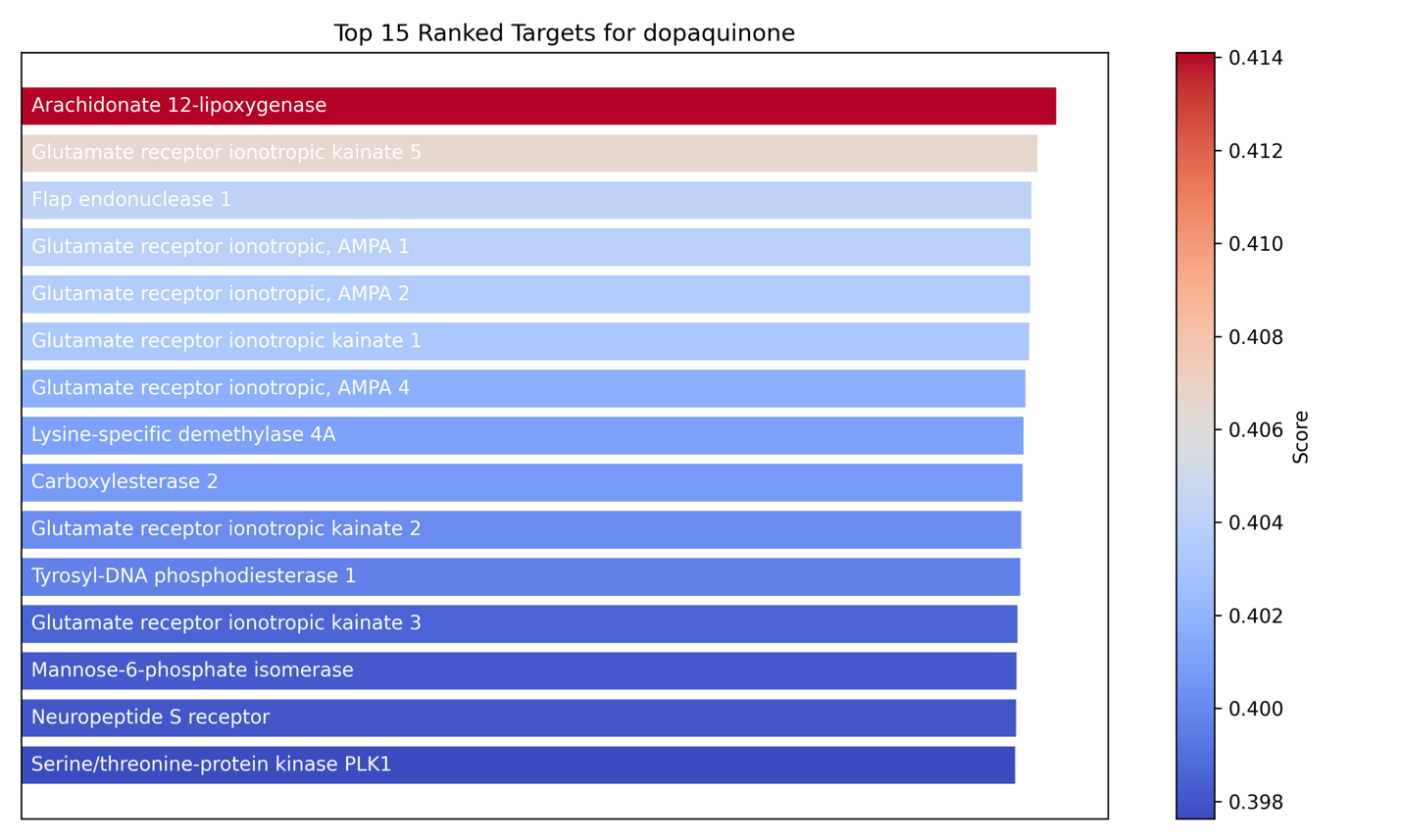

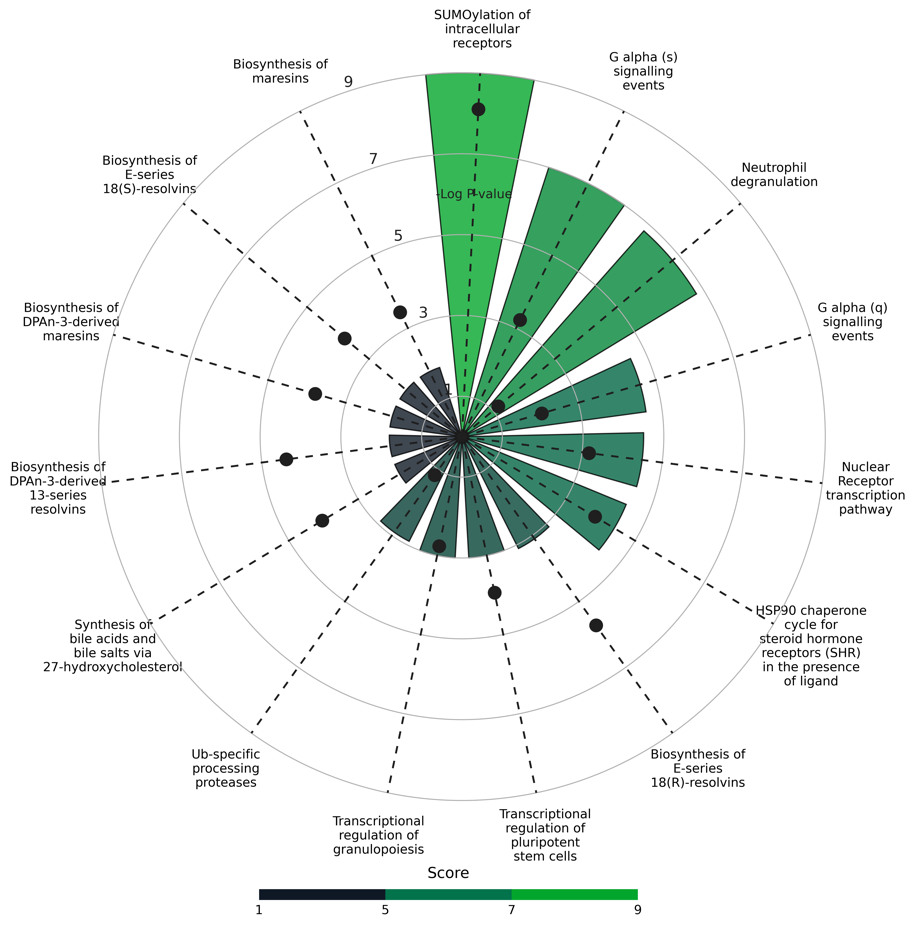

Figure S11. **Top 15 ranked targets predicted and mapped pathways for dopaquinone.**

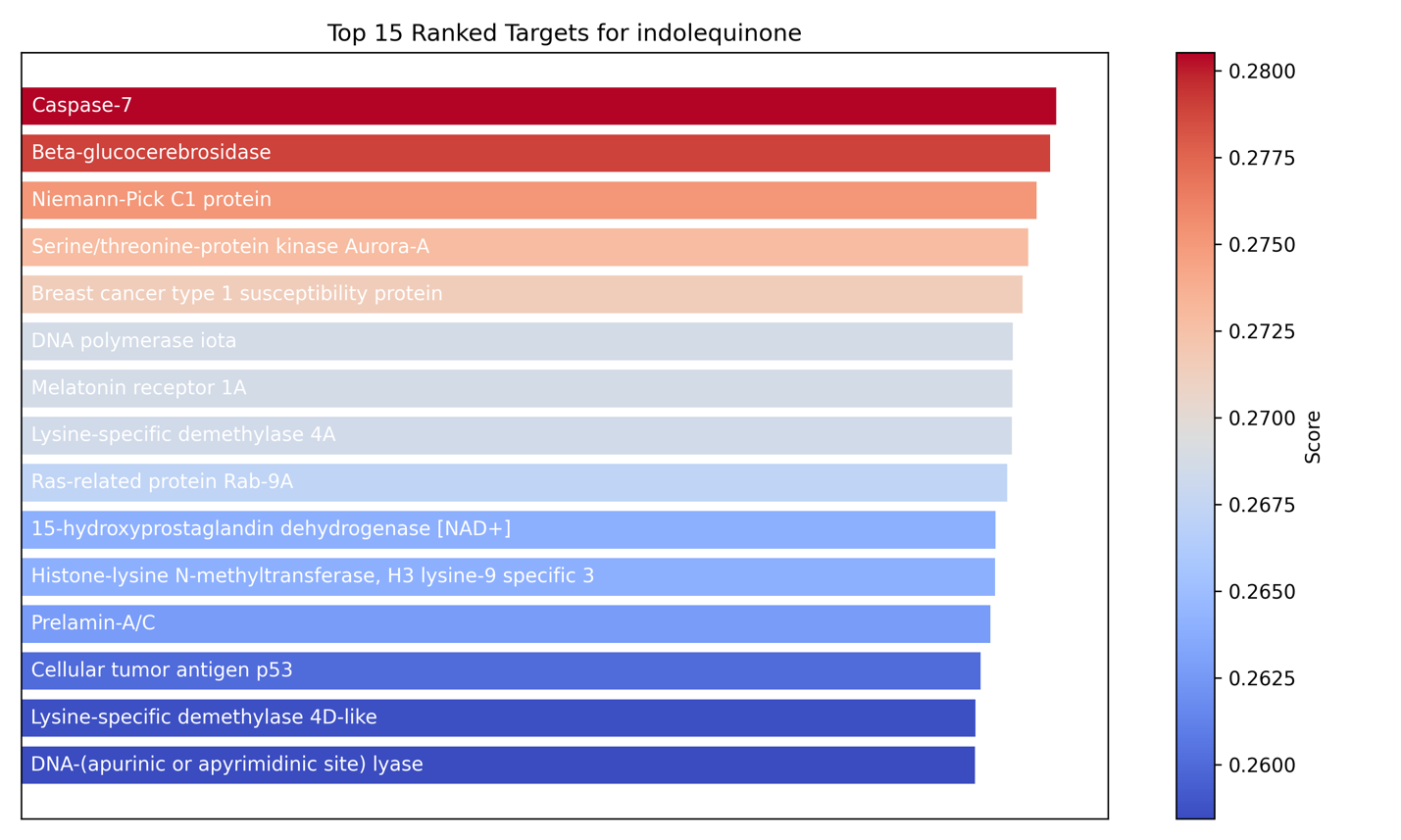

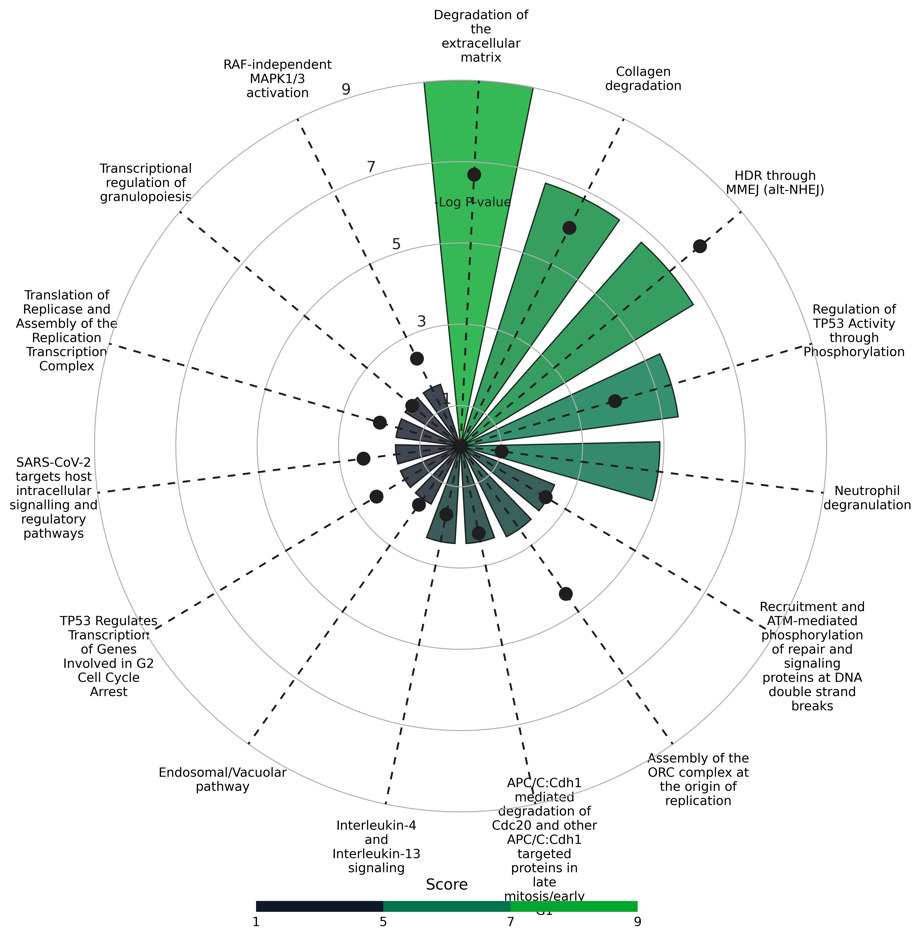

Figure S12. **Top 15 ranked targets predicted and mapped pathways for indolequinone.**

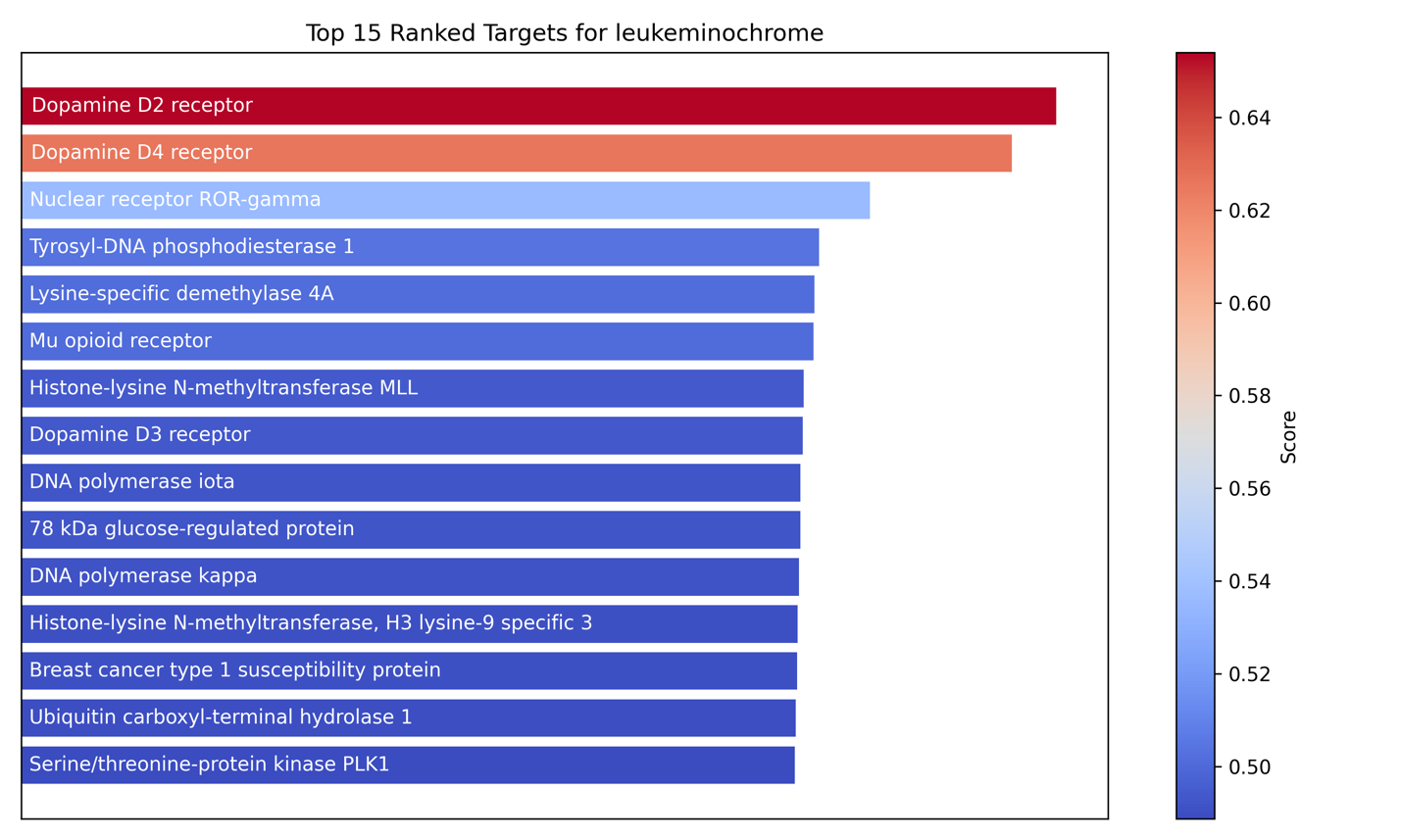

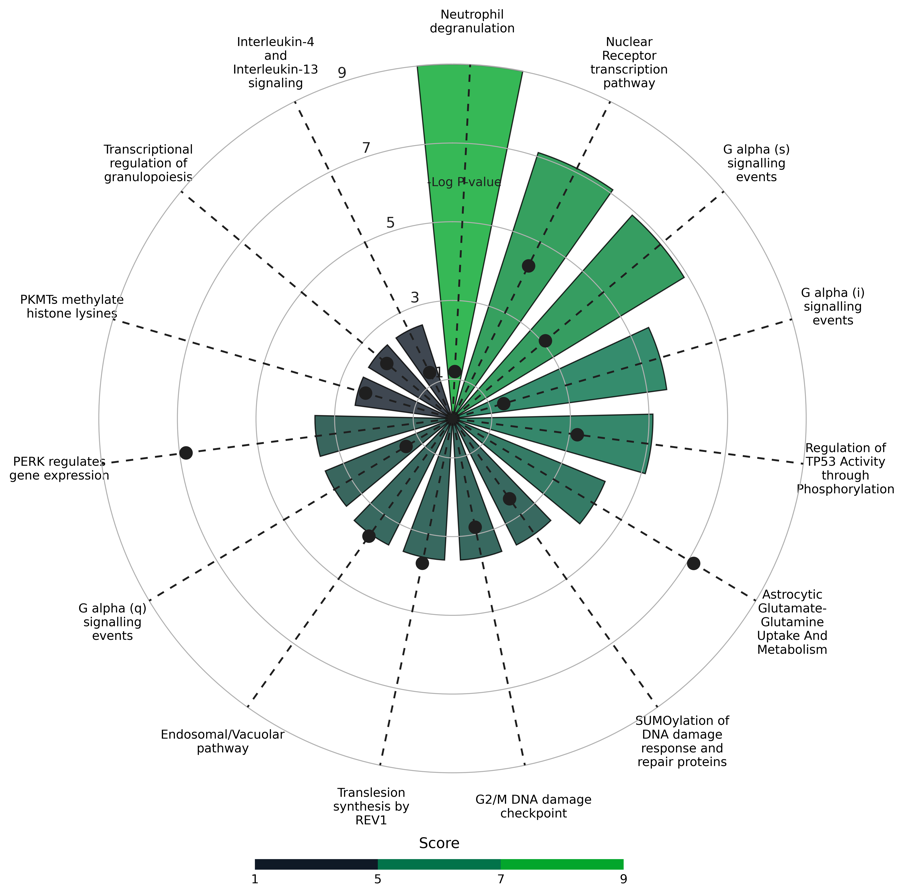

Figure S13. **Top 15 ranked targets predicted and mapped pathways for leukeminochrome.**

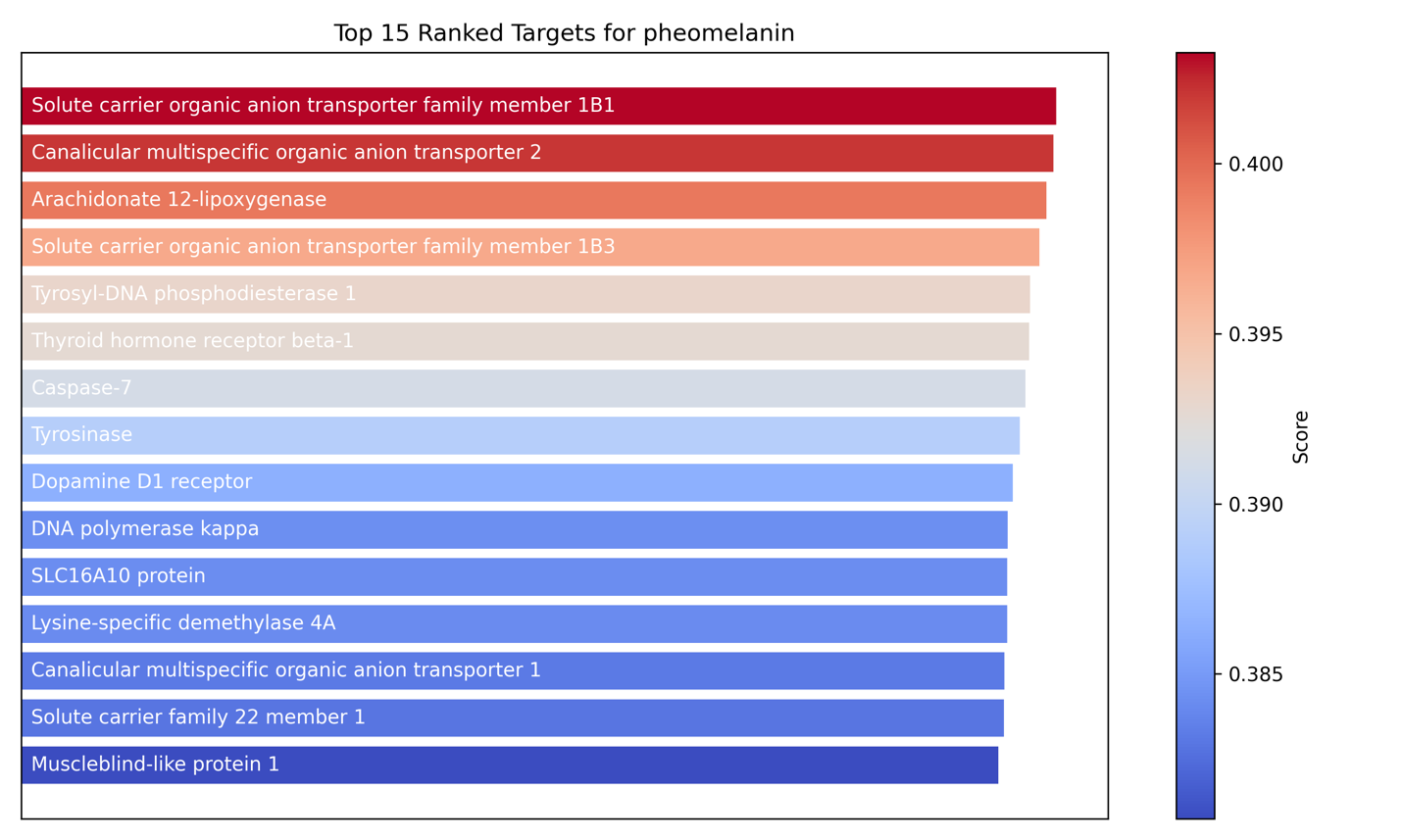

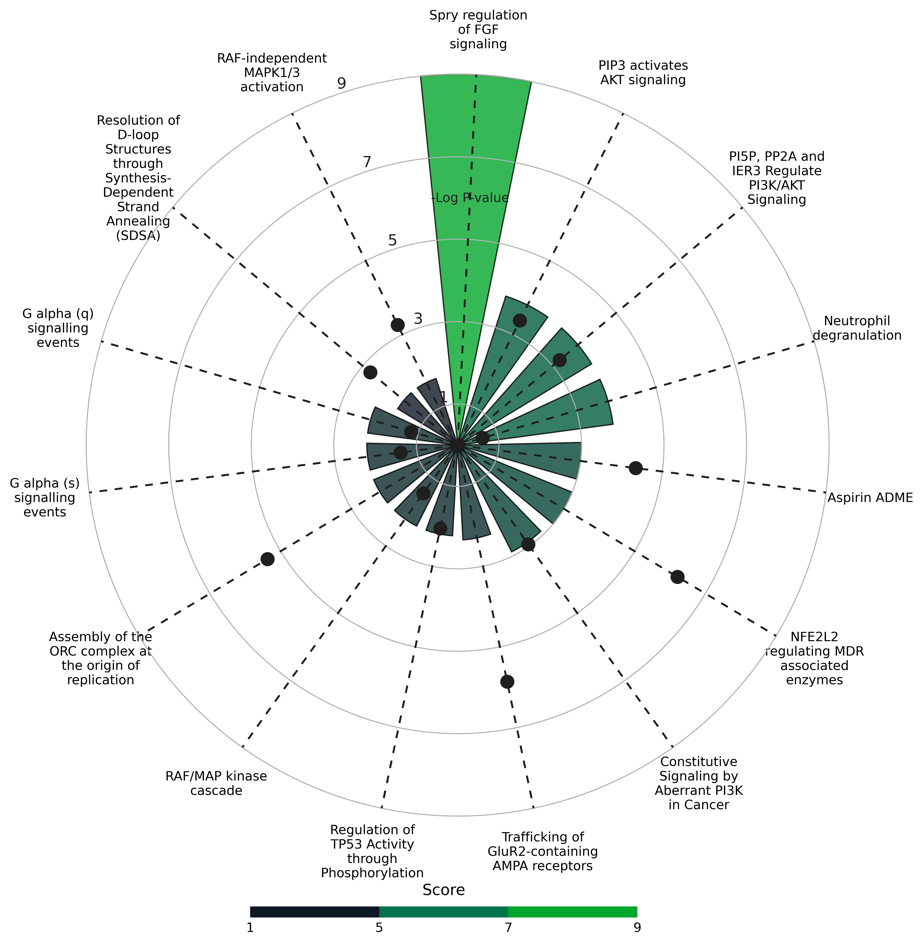

Figure S14. **Top 15 ranked targets predicted and mapped pathways for pheomelanin monomer**
